# Transposable elements are vectors of recurrent transgenerational epigenetic inheritance in nature

**DOI:** 10.1101/2024.09.20.614076

**Authors:** Pierre Baduel, Louna De Oliveira, Erwann Caillieux, Grégoire Bohl-Viallefond, Ciana Xu, Mounia El Messaoudi, Matteo Barois, Vipin Singh, Alexis Sarazin, Felipe K. Teixeira, Martine Boccara, Elodie Gilbault, Antoine de France, Leandro Quadrana, Olivier Loudet, Vincent Colot

## Abstract

Loss of DNA methylation over transposable elements (TEs) can affect neighboring genes and be epigenetically inherited in plants, yet the determinants and significance of this additional system of inheritance are unknown. Here, we demonstrate at thousands of TE loci across the Arabidopsis thaliana genome, that experimentally-induced hypomethylation can be transmitted transgenerationally and reveal the role of small RNAs derived from related copies in counteracting this transmission. Using data from >700 strains collected worldwide, we uncover natural hypomethylation at hundreds of the same TE loci, often situated near stress-responsive genes. Like their experimental counterparts, most natural epivariants we tested can be inherited without DNA sequence changes and are therefore bona fide epialleles, although genetic factors modulate their recurrence or persistence. Crucially, we demonstrate that TE-mediated epiallelic variation associated with differential gene expression is generally causal and may be target of selection in specific environments, thus establishing its importance in nature.

## Main Text

Transposable elements (TEs) are ubiquitous components of eukaryotic genomes and most TEs are ancestral copies that have long lost their ability to transpose(*1*). In plants and mammals, TEs are typically targeted by DNA methylation and loss of this epigenetic modification can result not only in increased TE mobilization, but also, whether or not TEs are transpositionally active, in chromosome rearrangements as well as altered expression of neighboring genes(*2*, *3*). Moreover, plants, in contrast to mammals(*4*, *5*), do not extensively reset DNA methylation across generations(*6–8*), opening up the possibility for the accidental loss of this modification to be inherited as epialleles, i.e. independently of DNA sequence changes. Indeed, a number of plant epialleles have been reported(*9–12*), some of which with striking consequences on fitness-related traits such as fruit development in oil palm(*13*) or the onset of flowering in *A. thaliana*(*14–16*). Crucially, most known epialleles involve ancestral, fixed or nearly-fixed TE insertions, thus raising the possibility that TE-mediated epiallelic variation has broad phenotypic impact in nature.

While undisputed in plants(*10*), TE-mediated transgenerational epigenetic inheritance (TEI) and its significance in nature remain poorly understood(*11*, *12*, *17*), in large part because of the difficulty to establish the independence of DNA methylation variation from DNA sequence polymorphisms(*18*). *Arabidopsis thaliana* offers a way forward however, thanks notably to several key resources and the deep understanding of the molecular players involved in DNA methylation of TEs developed in this model species. In particular, the Snf2-like chromatin remodeler DDM1 (DECREASE IN DNA METHYLATION 1), ortholog to human LSH/HELLS, was shown to be pivotal for the maintenance of methylation at the CG, CHG and CHH (where H=A,C,T) sites of TEs by allowing DNA methyltransferases access to heterochromatin(*16*, *19*–*22*), and crucially, *ddm1*-induced loss of DNA methylation can be stably inherited in the absence of the mutation(*23*). On the basis of this property, which most likely results from the disruption of the interplay between DDM1 and histone variants(*24*), a population of isogenic *ddm1*-derived epigenetic recombinant inbred lines (epiRILs) was generated that uncovered a large reservoir of potential epialleles affecting complex traits(*25–27*).

Here, we used this experimental epiRIL population to investigate the inheritance of DNA hypomethylation at thousands of TE loci across the genome and to characterize the molecular underpinnings of the wide differences observed. We then combined the knowledge gained in the epiRILs on TE-mediated epiallelic variation with genome sequence, DNA methylome, transcriptome, phenotypic and geo-bioclimatic data available for >700 *A. thaliana* strains from across the world(*28–31*), to interrogate the extent, dynamics, and implications of TE-mediated TEI in nature.

### Inheritance patterns of *ddm1*-induced hypomethylation differ markedly between TEs

To elucidate the molecular underpinnings of TE-mediated TEI in *A. thaliana*, we first obtained DNA methylome data at single nucleotide resolution for 120 *ddm1*-derived epiRILs(*25*) taken at generation F9 as well as for the *ddm1* and WT parental lines. The two symmetrical sites CG and CWG (where W=A or T) offer the most robust estimates of methylation levels and we identified 25,462 regions with markedly reduced methylation at these sites in *ddm1* (Fig. S1-2, Dataset S1-2). Using stringent criteria, including the presence of at least 5 differentially methylated CGs within a given differentially methylated region (DMR), we retained 7,023 DMRs (Table S1), hereafter designated as TE-DMRs, that are fully contained within TE annotations. Importantly, TE-DMRs do not belong to TE copies that are mobile in the epiRILs(*32*), thus alleviating the risk of mismapping of sequence reads in this setting. To determine the parental origin of each TE-DMR in the epiRILs, we “epihaplotyped” them (Fig. 1a, S2e, S3) using as genetic markers an independent set of 1,092 DMRs that are regularly spaced along the five chromosomes and which segregate the two parental methylation states in a manner compatible with Mendelian inheritance (Dataset S3). In a last step, we attributed for each TE-DMR one of three possible DNA methylation states in the epiRILs: WT-like, *ddm1*-like or intermediate. Specifically, we compared the average CG methylation level in each epiRIL with that in the two parental lines, as CG sites exhibit the largest differences of methylation between WT and *ddm1* (Fig. S1c).

**Figure 1.**
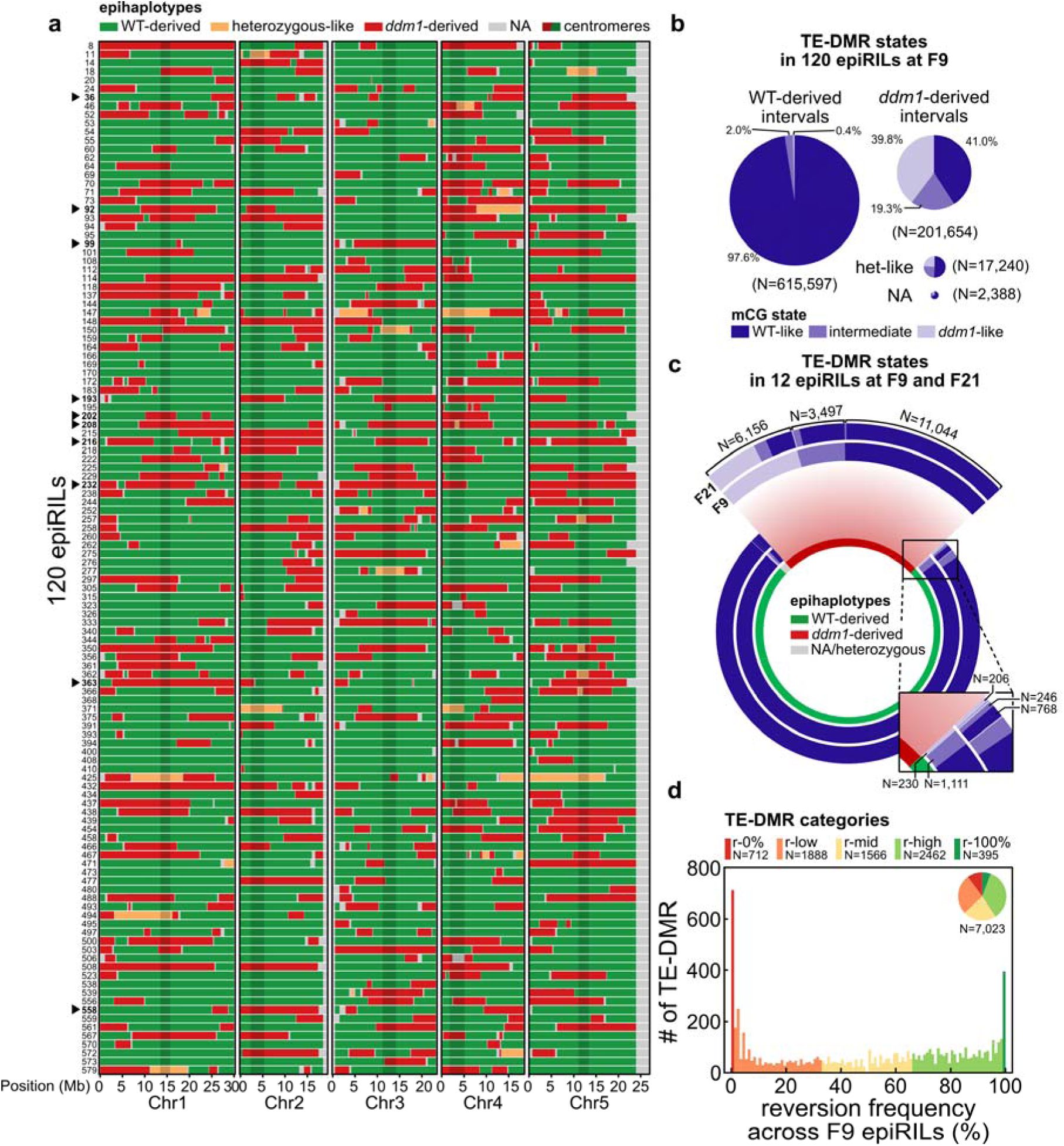
Inheritance patterns of *ddm1*-induced hypomethylation. (a) Epihaplotypic map of 120 F9 epiRILs. Centromeric regions are shaded. EpiRIL ID number is indicated on the left. (b) Distribution of methylation states (*ddm1-*like, intermediate, or WT-like mCG levels) of TE-DMRs within WT-derived or *ddm1*-derived intervals in the 120 F9 epiRILs. Black triangles point to the 10 contrasted epiRILs selected for RNA-seq. (c) Distribution of methylation states in the 12 epiRILs with F9 and F21 data. (d) Distribution of reversion frequencies measured at F9. Colors indicate the five categories of TE-DMRs we defined based on reversion frequency (r-0% to r-100%) and the pie chart indicates their relative proportion.

As previously reported using low-resolution DNA methylation data(*25*, *26*, *33*), while WT methylation is almost systematically inherited in the epiRILs (Fig. 1b, see Supplementary Note 1 for exceptions), this is not the case for *ddm1*-induced hypomethylation. Indeed, the vast majority of *ddm1*-derived TE-DMRs (N=6,311 or 89.1%) exhibit at least one instance of WT-like or intermediate DNA methylation in the epiRILs, indicative of full or partial reversion, respectively (Fig. 1b). Moreover, DNA methylome data produced from a random subset of 12 epiRILs propagated to generation F21 revealed that once restored in F9, WT-like DNA methylation is almost always (99%) stably transmitted to F21 (Fig. 1c) and that intermediate DNA methylation at F9 usually translates into WT-like levels by F21 (Fig. 1c), in agreement with the stability of WT methylation and the known progressivity of reversion(*33*), respectively (see Supplementary Note 2).

Reversion is conspicuous but far from systematic at generation F9, except for a small number (395) of TE-DMRs and we defined five categories of TE-DMRs to account for the wide range of reversion frequency at this generation: r-0%, r-low, r-mid, r-high, and r-100% (Fig. 1d). However, the observation that some r-0% TE-DMRs have reverted by generation F21 (Fig. 1b-c, S1e) suggests that *ddm1*-induced hypomethylation may not be inherited indefinitely in the absence of selection even at TE-DMRs of the most stable category.

### RdDM-mediated reversion is driven by non-allelic interactions between related TEs

Previous work established that the RNA-directed DNA methylation (RdDM) pathway, which involves 23-24nt small RNAs(*19*), is required for reversion(*33*). To investigate further the molecular underpinnings of reversion, we sequenced small RNAs from the two parental lines and 10 epiRILs. Consistent with earlier findings(*33*), the abundance of 23-24nt sRNAs in *ddm1* correlates positively with reversion (Fig. 2a, S4a), as a result of the drastic loss of 23-24nt sRNAs matching TE-DMRs that belong to the r-0% or r-low categories in this mutant background (Fig. 2a, S4b). Moreover, our new analysis reveals that this positive correlation is strongest for multiple-mapping 23-24nt sRNAs, which correlate also strongly with mCHH levels in *ddm1* (Fig. 2b, S4c). Given that the residual CHH methylation present in *ddm1* depends almost entirely on RdDM(*22*, *33*), the last two findings suggest a predominant role of multiple-mapping 23-24nt sRNAs in guiding RdDM activity and thus reversion. Consistent with this interpretation, reversion often affects related TE-DMRs together in the epiRILs (Fig. 2c) and it is more strongly associated with a higher abundance of multiple-than of single-mapping 23-24nt sRNAs not only in *ddm1* (Fig. 2a, S4a) but also in the epiRILs (Fig. 2d). Indeed, reversion is not accompanied by the restoration of WT levels of single-mapping 23-24nt sRNAs in some instances (Fig. S4d-e).

**Figure 2.**
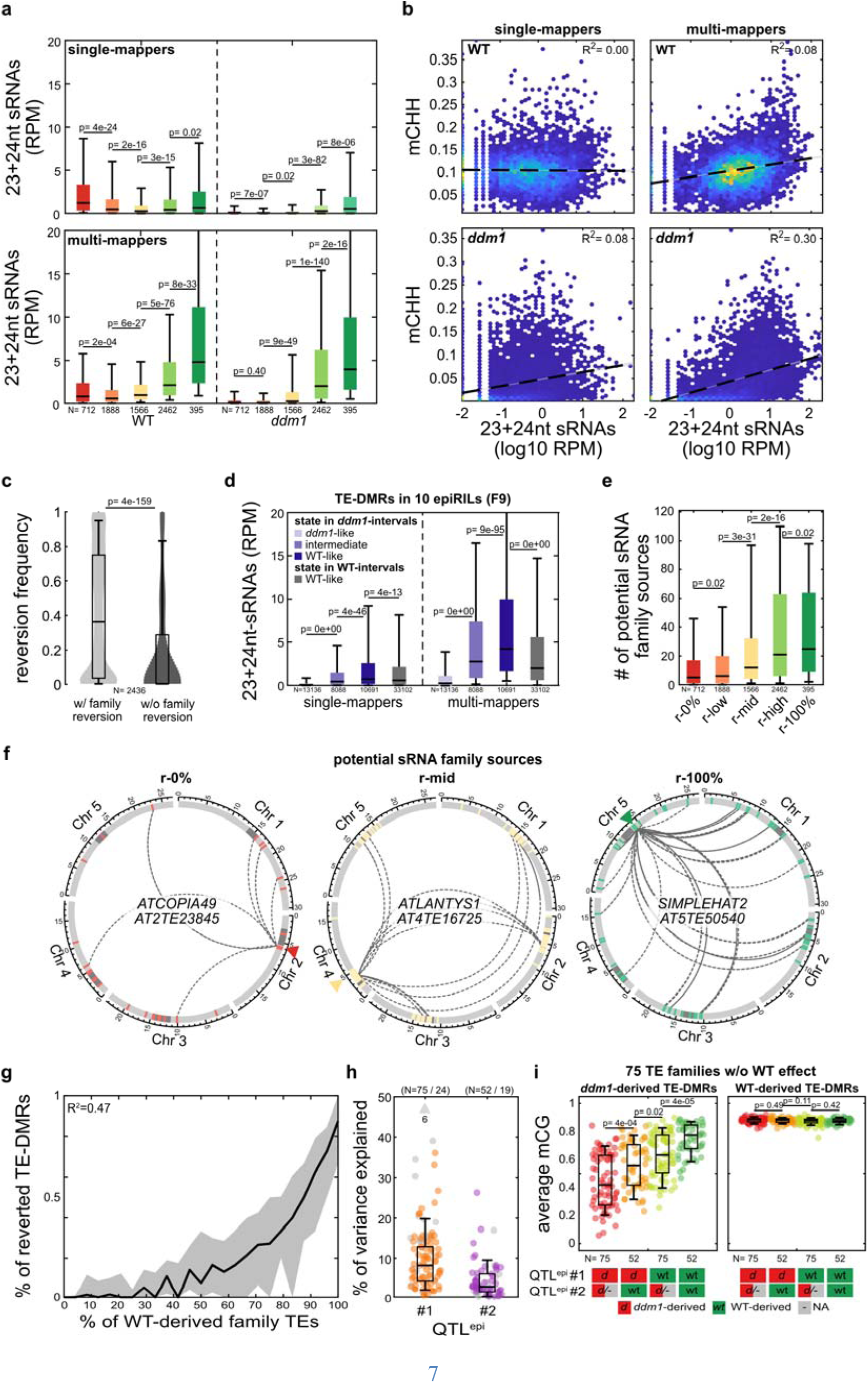
Reversion is driven mainly by *trans*-acting sRNAs. (a) Levels of 23-24nt single- (top) and multi-mapping (bottom) sRNAs in WT and *ddm1* over the five TE-DMR reversion categories. (b) Levels of mCHH vs levels of 23-24nt single- (left) and multi-mapping (right) sRNAs in WT and *ddm1* (log10 RPM). The coefficient of correlation (R^2^) of the linear regression (dotted line) is indicated. (c) Reversion frequencies of TE-DMRs in relation to whether or not the epiRILs exhibit reversion for at least one other TE-DMR of the same TE-family. (d) Levels of 23-24nt single- and multi-mapping sRNAs over TE-DMRs in 10 sequenced F9 epiRILs, depending on their parental origin and DNA methylation state. Only TE-DMRs that reverted in at least one of the 10 epiRILs were considered. (e) Number of potential 23-24nt sRNA sources belonging to the same TE family as the focal TE-DMRs, for each of the five reversion categories. (f) Genomic localization of three representative r-0%, r-mid, and r-100% TE-DMRs (indicated by triangles) and of related TE copies, including those that are potential sources of 23-24nt sRNAs (indicated by arcs). (g) Percentage of reverted TE-DMRs per TE-family relative to the proportion of related TEs that are WT-derived. (h) Percentage of variance explained iteratively by the first and the second QTL^epi^ in the average mCG levels of *ddm1*-derived, non-systematically reverting TE-DMRs within a given TE family. Values for the 24 TE families for which the QTL^epi^ affects the methylation status of WT-derived TE-DMRs are shown in gray, with outliers and their number indicated with a triangle. (i) Average mCG levels of *ddm1-* or WT-derived TE-DMRs per TE family in relation to the epihaplotype at the single or two major QTL^epi^ #1 or #2. Relations are only shown for the 75 TE families for which the QTL^epi^ do not affect the methylation status of WT-derived TE-DMRs.

To identify the potential sources of *trans*-acting 23-24nt sRNAs that guide reversion, and because RdDM can be driven by 23-24nt sRNAs with imperfect homology(*34*), we examined all 23-24nt sRNAs that can be aligned with up to one mismatch to target TE-DMRs. As expected, most (92%) of these sRNAs match perfectly to TEs that belong to the same TE family or superfamily as the target TE-DMR (Fig. S5a) and unsurprisingly, their number also correlates positively with reversion (Fig. 2e-f, S5b-c). Moreover, the proportion of TE-DMRs that belong to the same TE family and have reverted in a given epiRIL correlates positively with the number of related TE copies inherited from WT in that epiRIL (Fig. 2g). This last observation indicates that WT-derived sources are the main drivers of reversion, in agreement with the fact that most 23-24nt sRNAs matching TE-DMRs of the r-0% to r-mid categories are lost in *ddm1*.

To determine if some TEs play predominant roles in reversion, we performed QTL mapping for each of the 105 TE families with a sufficient number of (≥10) non-systematically reverting TE-DMRs, taking as a trait the average CG methylation of the TE-DMRs inherited from *ddm1*. Using this approach, we identified the presence of one or at most two major QTL^epi^ for almost all (99) TE families. QTL^epi^ intervals tend to overlap with TE-rich pericentromeres (Fig. S5d) and the WT-derived epigenotype in each case explains on average 13% of variation in family-wide reversion (Fig. 2h-i, S5e-f). Moreover, almost all (92%) QTL^epi^ encompass one or several of the potential sources of *trans-*acting sRNAs (Fig. S5d), which suggest that they are causal. Conversely, the *ddm1*-derived QTL^epi^ do not impact WT-derived TE-DMRs, except in rare cases that mainly concern TE families with known *trans*-demethylation activity (*VANDAL* and *ATENSPM* families, Fig. S5d-e, see Supplementary Note 1).

An important implication from the QTL mapping results is that the hypomethylated epiallelic state of non-systematically reverting TE-DMRs should be stable in epiRILs where the main sources of *trans*-acting 23-24nt sRNAs also derive from *ddm1*. To verify that this is the case, we grew >75 siblings of two randomly chosen F9 epiRILs (#92 and #216) that still harbor the hypomethylated epiallele at some non-systematically reverting TE-DMRs. We selected five TE-DMRs that belong to distinct TE families and have distinct reversion frequencies for analysis. Importantly, three TE-DMRs are members of TE families for which we identified at least one QTL^epi^ (Fig. S5d) and hypomethylation co-segregates with the *ddm1*-derived top QTL^epi^ in each case (Fig. S5g). Using targeted DNA methylation analysis, we detected zero or one reversion events among the F9 siblings, even for the two r-high TE-DMRs (Fig. S5f). These results indicate that hypomethylation can be stably transmitted in the absence of the sources of matching small RNAs that guide reversion. In other words, the wide differences in reversion frequency observed between TE-DMRs at generation F9 are determined primarily in *trans* rather than in *cis*.

### Transcriptional activity favors the inheritance but not the spontaneous occurrence of TE hypomethylation

Although reversion is usually concerted at the TE-family level (Fig. 2d), 369 TE-DMRs remained in the hypomethylated epiallelic state in epiRILs where over >75% of related TE-DMRs have reverted (Fig. S6a). Such “stray” TE-DMRs tend to have higher levels of sequence divergence with their closest relative and fewer close relatives altogether (Fig. 3a-b), in agreement with the known reduction of RdDM efficiency when source and target diverge in sequence(*35*). Furthermore, stray TE-DMRs are enriched at or near RNA Pol II transcription units that are activated in *ddm1* (EPigenetically Induced Consensus tAg clusTers or EPICATs(*36*); Fig. 3c-d, S6b-c, Supplementary Note 3), in keeping with the known inhibitory effect of Pol II transcription on RdDM(*37–39*).

**Figure 3.**
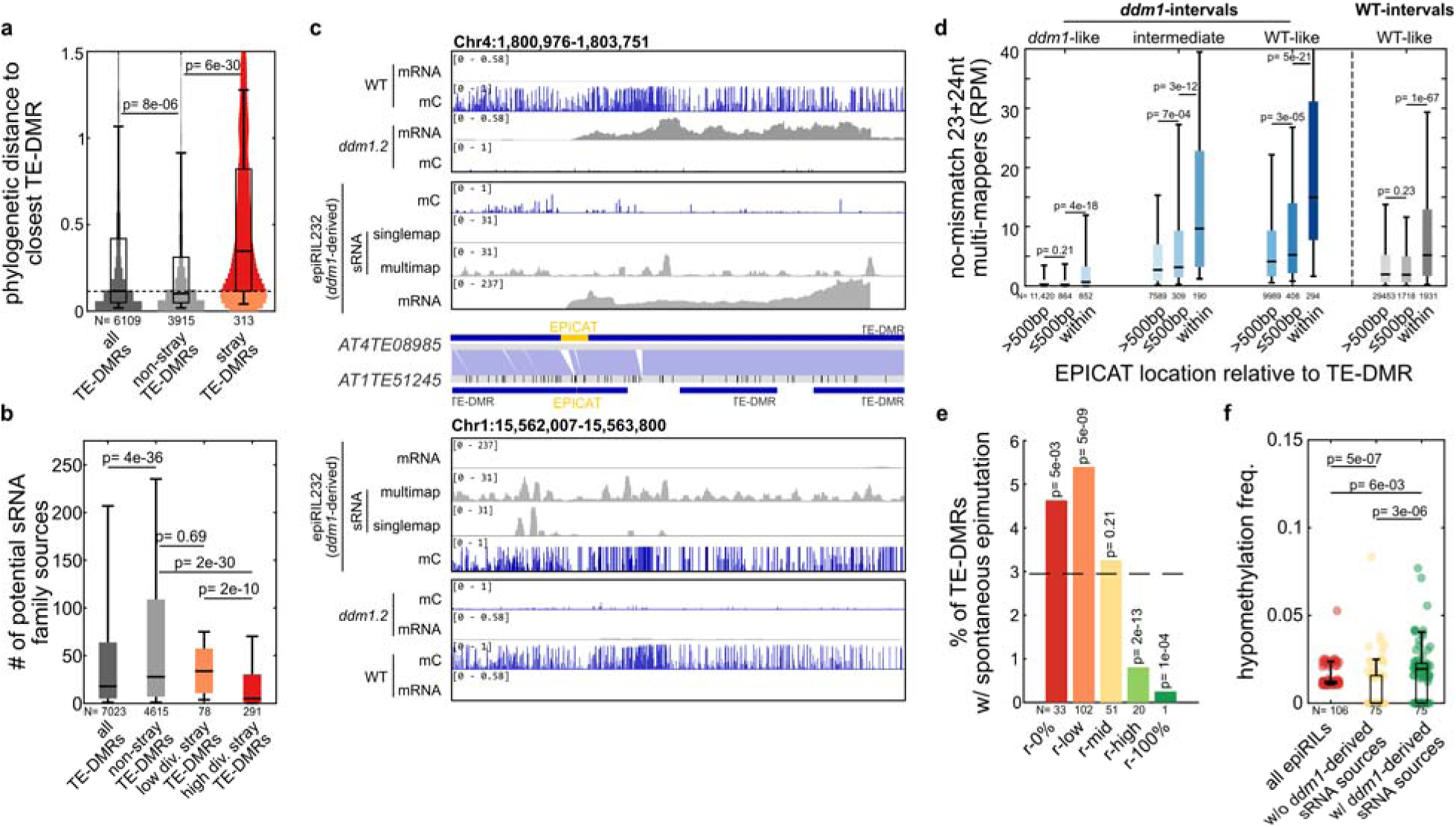
Inheritance and spontaneous occurrence of TE hypomethylation is modulated by two main antagonistic features. (a) Phylogenetic distance to closest TE-DMR for all, stray, and non-stray TE-DMRs. Stray TE-DMRs with distance lower and higher than the median are highlighted in orange (“low div.”) and red (“high div.”), respectively. (b) Number of potential 23-24 nt sRNA sources belonging to the same TE family for all, non-stray, “low div.” stray, and “high div.” stray TE-DMRs. (c) Levels of mRNAs(*36*), DNA methylation and 23-24 nt sRNAs, over a stray TE-DMR (*AT4TE08985*) that contains a long and strong *ddm1* EPICAT and which does not revert in epiRIL232, despite high levels of multi-mapping sRNAs and the presence in the same epiRIL of a closely related TE-DMR (*AT1TTE51245*) that has reverted. (d) Levels of multi-mapping 23-24nt sRNAs over TE-DMRs in 10 sequenced epiRILs (F9), in relation to their methylation state when they are WT- or *ddm1*-derived and as a function of their distance relative to the nearest EPICAT. (e) Percentage across the five reversion categories of TE-DMRs with spontaneous *ddm1*-like hypomethylation in the 120 epiRILs and mostly non-overlapping 169 epiRILs from Zhang et al.(*40*). The p-values indicate the cumulative binomial probability of observing similar or more extreme representation from random sampling (dashed line). (f) Frequency of spontaneous *ddm1*-like hypomethylation among all 120 epiRILs, only those where no related TE-DMR is *ddm1*-derived, or only those where at least one related TE-DMR is *ddm1*-derived. The presence of outliers and their numbers are indicated by a gray triangle.

To determine whether RdDM targeting may also affect the spontaneous occurrence of the hypomethylated state of TE-DMRs, we considered the 106 TE-DMRs with severe loss of parental WT methylation in one or two epiRILs only and that do not belong to TE families with known *trans*-demethylation activity (Table S4). To increase our sample size, we exploited methylome data independently obtained at generation F9 for a minimally overlapping set of 169 epiRILs(*40*) and identified in this set another 114 TE-DMRs with similar spontaneous *ddm1*-like hypomethylation (Table S4). The majority (65%) of the 207 TE-DMRs we identified in total belong to the r-0% and r-low categories (Fig. 3e) and remarkably, spontaneous hypomethylation is far less frequent (2.3-fold reduction) when all rather than some or none of the potential sources of *trans*-acting 23-24nt sRNAs are WT-derived (Fig. 3f). Thus, we conclude that RdDM not only limits the inheritance, but also the spontaneous occurrence of epiallelic variation at TE-DMRs. In contrast, the presence of EPICATs does not appear to have any detectable impact (Fig. S6d).

### TE epivariants are common in nature and most are recurrent, *bona fide* epialleles

Having determined the potential for epiallelic variation at thousands of TEs across the genome using the epiRILs, we then asked how this potential translates in nature. To this end, we leveraged WGBS-seq data available from leaf tissues for 720 natural strains(*29*) and calculated for each of the 7023 experimentally-defined TE-DMRs its average CG methylation level in each of the strains where it is present(*41*, *42*). Although TE-DMRs are generally part of TEs that are molecular fossils(*42–44*) and are thus fixed or nearly so at the species level (Fig. S7a-b), we retained for analysis only about half of them in any given strain because of the stringent filtering we applied to ensure robust measures of DNA methylation (Fig. S7c, Dataset S5).

In keeping with heavy and co-extensive DNA methylation being generally the default state for TEs in *A. thaliana*(*29, 45*), the vast majority (85%) of TE-DMRs have high average CG methylation across all informative strains (Fig. 4a). Nevertheless, 1,068 TE-DMRs exhibit severe, *ddm1*-like CG hypomethylation in at least one strain (Table S5). Most of these natural epivariants likely result from rare and rapid loss of DNA methylation, given that they usually have low population frequencies (Fig. 4a; Fig. S7d) and that intermediate levels of average CG methylation are seldom observed. Importantly, DNA methylation loss extends to CHG and CHH sites, as in *ddm1* (Fig. S7e,f), although the exact boundaries of this loss may differ (e.g. Fig. 4b). In addition, long-read genome sequencing of 20 genetically diverse strains (Table S6) indicates that the mismapping issues typically generated by the presence of related TEs in non-reference strains(*46*) affects only a small fraction (<15%) of the natural epivariants we called in each strain, thanks to our stringent filtering (Fig. 4c, see Supplementary Note 4).

**Figure 4.**
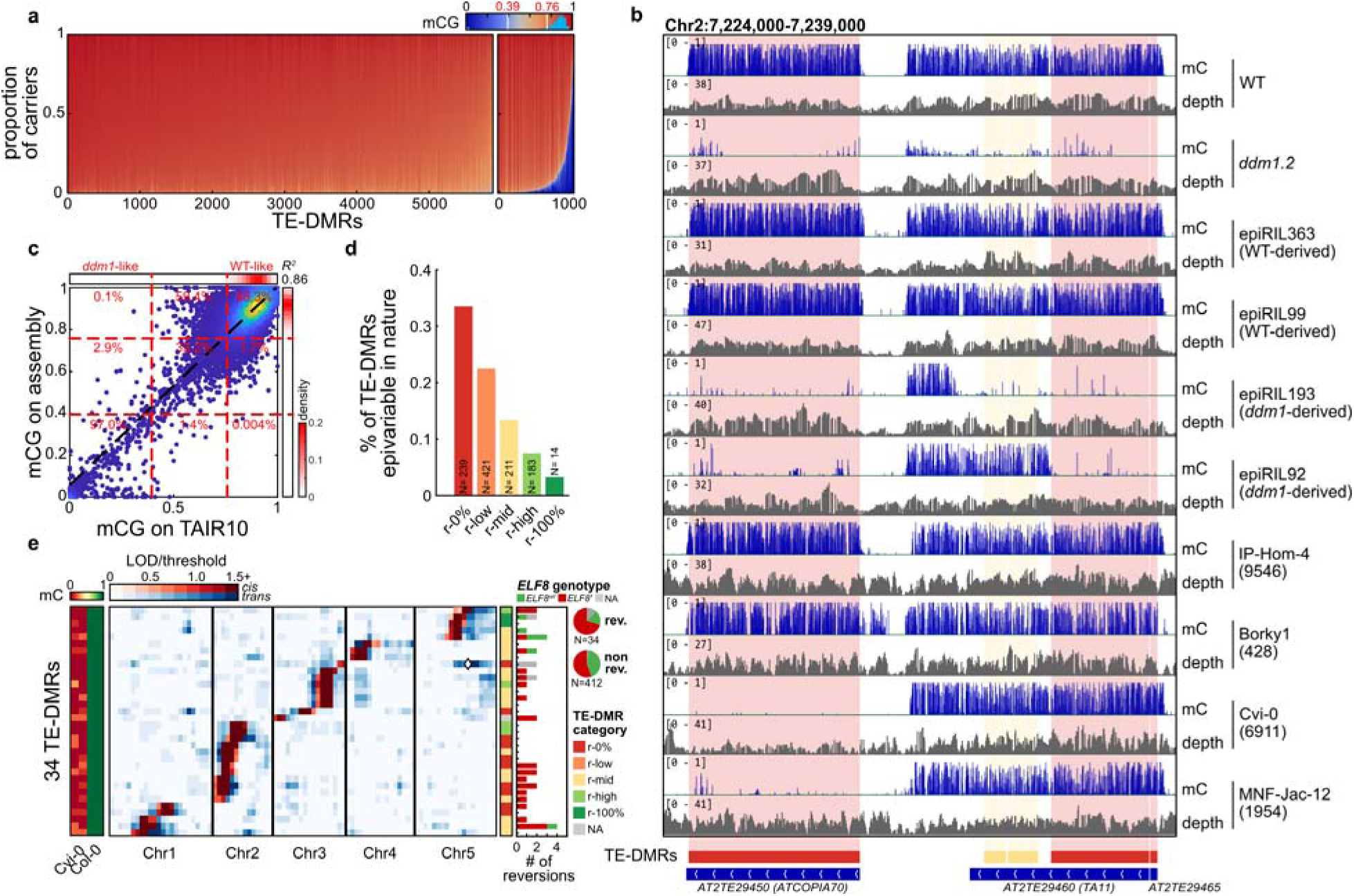
TE epivariants are recurrent in nature. (a) Distribution of average mCG levels over TE-DMRs in 720 natural strains. TE-DMRs with mCG under 0.39 were considered *ddm1*-like and over 0.76 as WT-like (density of average mCG levels is represented in cyan over the colorbar). (b) BS-seq mC levels and coverage over three TE-DMRs in the WT and *ddm1* parents, four epihaplotypically contrasted epiRILs and four natural strains in which two exhibit epivariation over one (*ATCOPIA70)* of the three TE-DMRs. (c) Comparison in 20 strains of average mCG levels measured by aligning BS-seq data on the *de novo* genome assembly vs on the TAIR10 reference genome assembly over TE-DMRs that are found at the reference location neither truncated nor missing in these 20 strains. The black dashed line represents linear regression. Red lines represent upper and lower thresholds for *ddm1*-like and WT-like methylation, respectively. Congruence is indicated by the percentage in each rectangle defined by these two thresholds. (d) Proportion of TE-DMRs with natural epivariation across the five reversion categories. (e) Heatmap of the mQTL mapping LOD scores in 36 Cvi-0 x Col-0 RILs for 34 TE-DMRs located across all five chromosomes. LOD score values are in shades of red or blue for associations in *cis* or trans, respectively. The unmethylated *ATCOPIA23* copy present in Cvi-0 (*42*) on Chr5 that is closely related to that present methylated on Chr3 in Col-0 (Fig. S8d), is indicated with a white diamond. The number of reversion events is represented for each TE-DMR as a bar colored by the *ELF8* genotype of the RILs in which they occurred. Frequency of reversion or stable inheritance by genotype are summarized in pie charts.

Naturally epivariable TE-DMRs are strongly skewed towards the r-0% and r-low categories (Fig. 4d), suggesting that the inheritance properties of *ddm1*-induced hypomethylated TE-DMRs extend to their natural counterparts. To confirm that this is the case and that natural epivariants are therefore *bona fide* epialleles, we first carried out genome-wide association studies (GWAS) of natural epivariation at each of the 490 TE-DMRs that have sufficient epivariant counts. Most significant associations with single nucleotide polymorphisms (SNPs) are in *cis* (within 20kb; Fig. S8a-b), yet none of these are systematic (Fig. S8c), thus ruling out a strict dependence of natural epivariation on neighboring SNPs.

To obtain direct evidence that polymorphisms in *trans* do not dictate epivariation either, we performed linkage mapping using the population of recombinant inbred lines (RILs) previously built between Cvi-0 and Col-0(*47*). This RIL population is particularly informative in this respect, as Cvi-0 harbors the second largest number of epivariants (113 in total, Fig. S7c) among natural strains and also because almost all (110) of the corresponding TE-DMRs are hypomethylated in at least one other strain (Table S7). Targeted (Fig. 4e) as well as whole-genome (Fig. S8e) segregation analyses indicate that the two parental DNA methylation states almost systematically segregate in the RILs together with flanking sequence markers, and thus independently from sequence variants in *trans* (see Supplementary Note 5 for the one exception). In addition, many of the Cvi-0-derived TE-DMRs have regained dense DNA methylation in at least one RIL (Fig. 4e, S8e), which confirms that they are also not strictly conditioned by polymorphisms in *cis*.

Epivariants present in two or more strains could reflect identity by descent or recurrent occurrence. To distinguish between these two possibilities, we considered the 780 TE-DMRs with several instances of epivariation in nature and calculated the genetic relatedness (kinship) of the corresponding strains. While almost all (96%) TE-DMRs show hypomethylation in at least two strains that are highly divergent (genome-wide kinship <0.25), only half of them do so in two or more closely related strains (genome-wide kinship >0.75; Fig. S9a). Furthermore, local haplotyping indicates that for the majority (66%) of TE-DMRs, epivariants are carried by at least two distinct alleles (Fig. S9b-d). Thus, the population frequencies of natural epivariants, are determined in large part by recurrence rather than long-term inheritance, consistent with their experimental epiallelic counterparts being not heritable indefinitely in the absence of selection.

### Natural epivariants are prevalent near genes

Unlike in the epiRILs, natural hypomethylated epivariants are generally few in any given strain (49 on average, Fig. 4b, S7c) and tend to be either tightly clustered (within 10kb) and correlated between strains, or else dispersed and uncorrelated (Fig. 5a, S9e-f). In addition, naturally epivariable TE-DMRs are usually less pericentromeric (Fig. 5b), hence closer to genes than the complete set of experimental TE-DMRs (Fig. 5c). Importantly, the population frequency of natural epivariants correlate positively with gene proximity, suggesting that it facilitates either their occurrence or inheritance, or both. Further supporting this interpretation, we identified through GWAS the gene *EARLY FLOWERING 8* (*ELF8*) as the single major genetic determinant of the prevalence of natural epivariation within strains (Fig. 5d-f). *ELF8* encodes a component of the highly conserved RNA polymerase-associated factor 1 (Paf-1) complex, and previous work showed that when marked by the histone H3 modification H3K4me2/me3(*48*), which in plants requires Paf1(*49*), genes hinder RdDM targeting of neighboring TEs. Consistent with these findings, strains with the non-reference, overexpressing allele of *ELF8* (hereafter designated *ELF8’*, see Fig. S10a-h and Supplementary Note 6 for functional analysis) tend to harbor a higher number of natural epivariants (Fig. 5e,g) and this trend is most pronounced for TE-DMRs that are located within 2kb of genes (Fig. S10i). However, examination of the Cvi-0 x Col-0 RILs, which segregate the two *ELF8* alleles, indicate that increased ELF8 activity is insufficient on its own to promote the occurrence of epivariants or to stabilize their inheritance (Fig. 4e, S10j-k, see Supplementary Note 7). Thus, additional factors are likely involved in the association observed in nature.

**Figure 5.**
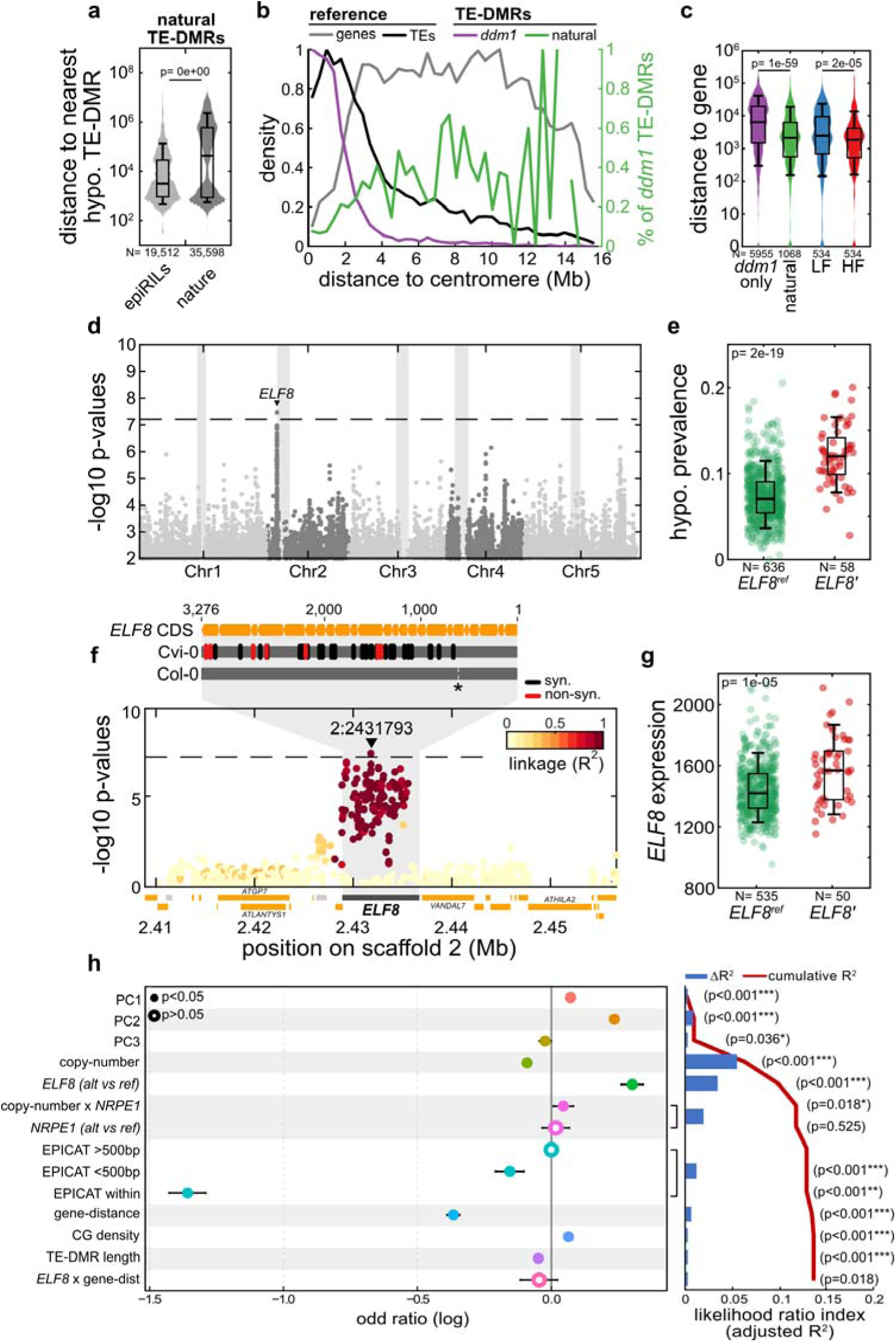
Natural epivariants are enriched near genes. (a) Distance between closest naturally epivariable TE-DMRs in the hypomethylated state in the epiRILs and in natural strains. (b) Metaplot of the density of TEs, genes, *ddm1*-induced TE-DMRs in relation to their distance from centromeres (left y-axis). The proportion of *ddm1*-induced TE-DMRs that are also epivariable in nature is represented in green (right y-axis). (c) Distance to the nearest gene (+1bp for log representation) of all *ddm1*-induced TE-DMRs, naturally epivariable TE-DMRs and those for which epivariation segregate at low or high frequency (LF, first two quartiles; HF, last two quartiles). (d) Manhattan-plot of p-values of GWAS of prevalence of natural epivariation by genome. (e) Prevalence of natural epivariation among strains carrying the reference *ELF8^ref^* or non-reference *ELF8’* alleles. (f) Close-up Manhattan-plot of GWAS p-values around *ELF8,* colored by linkage with the leading SNP (2:2431793). Coding-sequence polymorphisms with the Cvi-0 *ELF8’* allele are indicated above. Asterisk marks a frame-shifting single-nucleotide deletion in the Col-0 TAIR10 sequence that is not detected in re-sequenced Col-0 *ELF8* cDNA (see Supplementary Note 6). (g) *ELF8* expression in strains carrying the *ELF8^ref^* or *ELF8’* alleles. (h) Marginal effects (log odd ratio) and likelihood ratio index (McFadden adjusted R^2^) of each explanatory variable in GLM of natural epivariation prevalence ranked by R^2^.

To explore what these additional factors might be, we used a generalized linear model (GLM) and performed a stepwise selection of ten parameters, including allelic variation at *ELF8* and distance to genes, on the basis of their added explanatory power. Results of the GLM indicate that TE copy-number and allelic variation at *ELF8* are the two main parameters, which explain 5.5% and 3.5% of the variance in the proportion of TE-DMRs that are hypomethylated in a given strain, respectively (Fig. 5h). The negative correlation with TE-copy number is consistent with RdDM strength increasing with TE copy-number(*50*, *51*) and the role of RdDM in limiting both the inheritance and spontaneous occurrence of hypomethylation at TE-DMRs in the epiRILs (Fig. 2-3, S5-6). Indeed, our GLM revealed that the weak allele of the RdDM gene *NRPE1*(*42, 52*) mitigates the negative effect of TE copy-number on epivariation. Moreover, it also confirmed that the impact of gene proximity is stronger in strains carrying the *ELF8’* rather than the *ELF8^ref^*allele (Fig. S10i). Together, these findings highlight the pivotal and antagonistic roles of RdDM and Paf1 activity in shaping the prevalence of epivariation in nature.

### Natural epialleles impact gene expression

To investigate the functional impact of natural epivariation, we first analyzed RNA-seq data available for >700 strains(*29*) and compared for each naturally epivariable TE-DMR the expression levels of the closest genes between strains with or without epivariation. Although we probed just one organ (leaf) and one condition (standard growth), we observed major (≥2-fold) gene expression changes in *cis* for 25% of TE-DMRs (Fig. S11a). These changes were strongly skewed towards upregulation, notably when the TE-DMRs overlap with genes or are located immediately upstream (≤500bp) of them (Fig. S11a-b). To distinguish between cause and consequence, without the possible confounding effects of DNA sequence variants, we produced RNA-seq data from the 10 epiRILs we used for the sRNA studies and asked whether the experimental counterparts of epivariants associated with changes in gene expression in nature cause similar changes in the epiRILs. Comparisons could be made for 99 TE-DMRs in total. For 58 of them the expression changes between epiRILs with contrasted epiallelic states were congruent, both in direction and amplitude, with those in nature (Fig. 6a, S11c) thus explaining 24% on average of the variation in expression levels observed between natural strains (Table S10). In addition, congruence concerns 30 of the 32 TE-DMRs located within or upstream (≤500bp) of genes (Fig. 6a, S11c) and for 14 of the 17 TE-DMRs with a reversion event in the 10 epiRILs sampled, WT expression is regained too (Fig. 6b, S11d). Thus, the majority of natural epiallelic variants associated with gene expression differences in *cis* are causal and causality is almost systematic for TE-DMRs within or upstream of genes.

**Figure 6.**
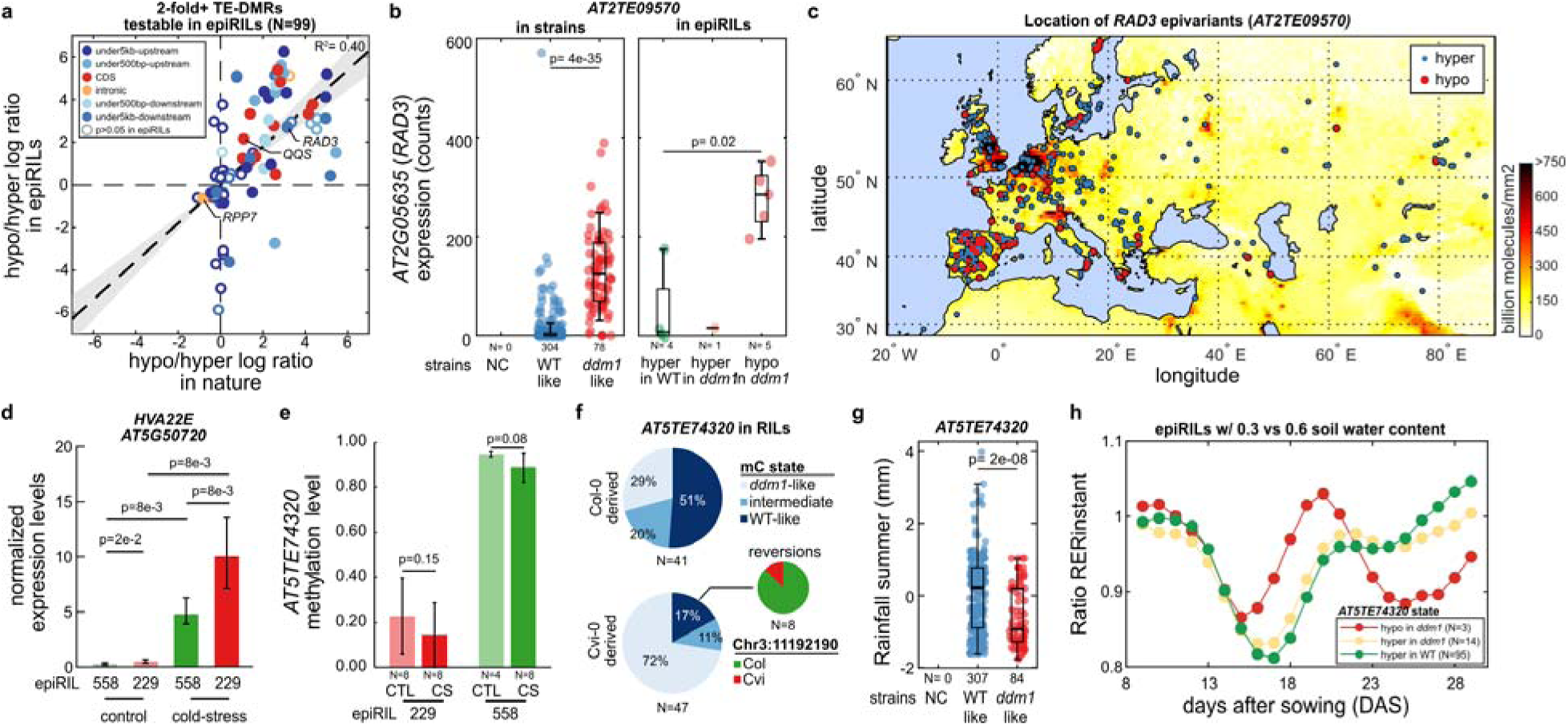
Natural epivariation has functional consequences. (a) Comparison of the hypo/hyper log_10_ ratios measured in the epiRILs and in nature for each of the 99 naturally epivariable TE-DMR with ≥2-fold gene expression differences in nature and for which the 10 epiRILs are informative. Position of TE-DMRs relative to the affected genes is indicated by a color code. (b) Comparison of *RAD3* expression level between strains with or without epivariation at the fixed *RAD3* TE-DMR (left panel) and between the 10 epiRILs according to the epiallelic state of the *RAD3* TE-DMR (right panel). (c) Localization of Eurasian *A. thaliana* strains with epivariation at the *RAD3* locus against NO_2_ atmospheric levels in June 2005 (AURA_NO2_M_2005-06-01 from neo.gsfc.nasa.gov) (d) Expression level of *HVA22E* measured in three independent pools of 8 F10 seedlings of each of the epiRIL229 and epiRIL558 (with and without *ddm1*-induced hypomethylation of the downstream TE-DMR, respectively) grown under control conditions or exposed for 24h to cold-stress. (e) DNA methylation levels at *AT5TE74320* TE-DMR measured in 4-8 F10 individuals of the epiRIL229 and epiRIL558 grown under control conditions or exposed to cold-stress for 24h. (f) Methylation states of *AT5TE74320* within 88 Cvi-0 x Col-0 RILs (*47*) depending on genotypes at both bordering markers (left two pie charts). Genotype at top marker within QTL^epi^ interval for 8 RILs with reversion (right pie chart). (g) Comparison of rainfall in summer (2001–2010) at the collection site of strains with or without epivariation at the fixed TE-DMR (NC=0) downstream of *HVA22E*. The presence of outliers and their numbers are indicated by a red triangle. (h) Comparison of the difference in daily relative growth rate (RER ratio) of epiRILs grown on the Phenoscope in mild-drought vs well-watered conditions depending on the epiallelic state of the *HVA22E* TE-DMR.

Many of the 60 genes for which we could prove causality have no known function and the phenotypic consequences of their misregulation remain to be determined. Among those with known function (Table S10), two were previously shown to be affected by the epigenetic state of the TE sequences they contain (*QQS*(*53*) and *RPP7*(*54*)). While the phenotypic impact of the epiallelic upregulation of *QQS* (Fig. S11e) remains to be determined, the inhibition of splicing caused by the loss of heterochromatin at the intronic TE present in *RPP7* (Fig. S11f), was shown to considerably reduce RPP7-mediated defence against pathogens(*55*). Thus, epiallelic variation at *RPP7* may be deleterious in natural settings, and consistent with this interpretation, we only observed it once among informative strains, despite the TE-DMR being of the r-low category. *RAD3* (*AT2G05635*), which encodes a DEAD helicase, also stands out because it lies within the major QTL^epi^ interval identified for primary root length in the epiRILs(*27*). Moreover, *RAD3* has been implicated in natural variation for root growth under conditions of acid mine drainage(*56*). Our transcriptomic analysis also shows that loss of DNA methylation at *RAD3* results in more abundant and longer transcripts (Fig. 6b, S12a) and is associated in the epiRILs with reduced growth of the primary root (Fig. S12b). Importantly, epivariants of *RAD3* are relatively frequent in nature in nature (95 strains) and biogeoclimatic data(*57*) indicate that they are underrepresented in locations with high NO_2_ emission levels in summer (Fig. 6c, S12c-d). As NO_2_ emissions correlate positively with levels of soil acidification(*58*), it is therefore tempting to speculate that hypomethylation of the *RAD3* TE-DMR is selected against in habitats with acidic soils. Thus, in addition to allelic variation, epiallelic variation at *RAD3* has phenotypic consequences that could be ecologically relevant.

### Natural epivariation may be selected to facilitate stress responses

Given that naturally epivariable TE-DMRs are particularly enriched near stress-responsive genes (Fig. S12e), we asked if epivariation could influence gene expression under stress conditions. For this purpose, we focused on the r-mid TE-DMR that is located ∼900bp downstream of the cold- and drought-stress responsive gene *HVA22E* (*AT5G50720*; Fig. S13a). In each of the two pairs of epiRILs that we investigated with contrasted parental epiallelic states at the TE-DMR, *HVA22E* expression is similarly low in the absence of stress, but its induction by cold is markedly higher (1.6 and 2.4-fold) when the TE-DMR is hypomethylated (Fig. 6d, S13b). Furthermore, cold treatment does not result in a loss of DNA methylation in the case of the WT-derived TE-DMR (Fig. 6e, S13c). Altogether, these results reveal that the hypomethylated epiallele at *HVA22E* enables plants to enhance their response to cold stress, without affecting their basal expression.

Hypomethylation at The TE-DMR downstream of *HVA22E* is quite common (24%) among natural strains, despite reversion being frequent (66%) in the epiRILs (Table S1), which suggest the presence of stabilizing genetic or genomic modifiers, or positive selection in nature. Although discriminating between these possibilities can be challenging, we exploited the fact that the TE-DMR is in the hypomethylated state in Cvi-0 to assess its stability in 88 Cvi-0 x Col-0 RILs. The Cvi-0 allele remained hypomethylated in 72% of the of the RILs that inherited it but regained high levels of DNA methylation in eight of them (Fig. 6f), seven of which harbor the Col-0 haplotype at the top marker within the QTL^epi^ interval we identified for this TE family in the epiRILs (Fig. S13d). These observations, together with the fact that the Col-0-derived allele loses DNA methylation in 29% of RILs indicate that the Cvi-0 genotype stabilizes hypomethylation at *HVA22E* and conversely, that the Col-0 QTL^epi^ has a strong destabilizing effect.

We observed that hypomethylation at *HVA22E* is preferentially found in habitats with high summer frost and low summer precipitation (Fig. 6g, S13e-f). Furthermore, analysis of phenotypic data compiled in the AraPheno database (https://arapheno.1001genomes.org/) indicates that strains with hypomethylation at the locus tend to have smaller stomata (Fig. S13f-g), a cell-type where *HVA22E* is strongly induced by drought(*59*). These observations prompted us to determine if epiallelic variation at *HVA22E* could impact the ability of plants to respond to drought. Specifically, we used a high-throughput phenotyping platform(*60*) to monitor the growth of 112 epiRILs subjected or not to mild drought, including three with a *ddm1*-derived epiallele and 14 where the *ddm1*-derived TE-DMR had reverted. While all epiRILs were equally affected by drought during the first seven days of growth under this condition, the epiRILs with the hypomethylated epiallele resumed growth faster than all of the other epiRILs (Fig. 6h, S13h), indicating that it could act as a selectable engine of enhanced response to stress.

## Discussion

By combining experimental and natural population epigenomics, we conducted a comprehensive study of the molecular underpinnings and biological significance of TE-mediated TEI in plants. Using the *ddm1*-derived epiRILs, we demonstrated that inheritance of hypomethylation at a given TE is determined in *trans*, because RdDM, which drives reversion, is guided by 23-24nt sRNAs that derive from related TE copies mainly. On the basis of this new knowledge obtained for thousands of TE loci across the genome, most of whom are ancestral, we then interrogated the presence and functional impact of natural epivariants with similar loss of DNA methylation over the same TE loci in nature. Despite stringent filtering and biases inherent to our study design (see Supplementary Note 8), we identified hundreds of TE loci with natural *ddm1*-like epivariation and showed that they tend to be located near stress-responsive genes. These natural epivariants reflect abrupt loss rather than gradual erosion of DNA methylation across generations and most of them are recurrent, *bona fide* epialleles that are relatively short-lived, like their experimental counterparts. Moreover, epiallelic variation at TE loci associated with changes in gene expression in *cis* is generally causal, especially when the TEs are located within or upstream of genes. Finally, we showed that effects can be more pronounced in response to stress or specific to it and that causal epialleles are not randomly distributed across habitats, suggesting selection is at play. Our study therefore provides a detailed understanding of the mechanisms responsible for TE-mediated TEI in plants and demonstrates the importance of this additional system of inheritance in nature.

The demonstration that sRNAs derived from related TEs play a prominent role in limiting TE-mediated TEI is in line with 23-24nt sRNAs being required in large amounts for efficient RdDM-targeting of transgenes(*37*, *61–64*), indicates that the determinants of epigenetic stability at a given TE locus are intrinsically polygenic. Thus, the genome could be viewed as an epigenetic biome in which the epiallelic properties of a given locus relies on interactions, additive or not, between multiple loci. Furthermore, the observation that pericentromeric TEs are the main source of *trans-*acting sRNAs and have little impact in *cis* is consistent with methylome remodeling data obtained in epigenetic F1 hybrids(*65*) and lends direct support to the proposition that these sRNAs mainly serve to guide DNA methylation of related copies located on chromosome arms(*66–69*). More generally, the finding that almost all TEs we analysed can sustain TEI in at least one epiRIL indicate that a one-to-one ratio of sources of 23-24nt sRNAs to targets is usually insufficient to direct RdDM to these, unlike in “classical” paramutation, which involves allelic or epiallelic interactions, such as at the *b* locus in maize(*70*). As a matter of fact, classical paramutation is rare in the F1 progeny of crosses between *A. thaliana* strains(*71*), which our analysis of the Cvi-0 x Col-0 RILs also indicate, and it may be the exception rather than the rule, as previously proposed based on more limited data (*37*, *72–74*).

Our analysis of natural epivariation revealed that while RdDM reduces the possibility of TE-mediated TEI, gene proximity and increased expression of *ELF8,* which encodes a component of the evolutionary conserved Paf1 complex, are the two main genetic factors that favor it. These findings are in line with results in *Schizosaccharomyces pombe* (*75, 76*) and *A. thaliana*(*48*) indicating respectively that Paf1 and the active histone marks it helps deposit inhibit small RNA-mediated epigenetic silencing. Thus, in addition to its many roles in facilitating Pol-II transcription(*49*), Paf1 may also help safeguard genes from the silencing of neighboring TEs.

We showed that natural epialleles can alter the expression of neighboring genes and that their effects may be exacerbated in response to stress. Furthermore, their presence does correlate with specific environments in some cases, in line with evidence indicating that despite their lower stability compared to DNA sequence variants, natural epialleles can be targets of selection (*27*, *77*). Given their recurrence and singular inheritance properties, these natural epialleles therefore may contribute uniquely to local adaptation, especially in cases where their effects on gene expression are minimal in the absence of stress. However, determining the extent of this contribution remains a major challenge, because of the difficulty in teasing apart the effects of epialleles and DNA sequence variants in natural populations. Moreover, a key open question is whether epiallelic variation may be facilitated or stabilized in specific environments, as evidence in support of such effects is still lacking (this study and(*78*)), despite well-documented effects of biotic or abiotic stresses on the heritable loss of methylation at individual cytosines(*79–82*). Whatever the case might be, our findings clearly establish that ancestral TEs are an important and unique source of recurrent heritable variation in plants, with broad implications for conservation efforts and breeding programs.

## Acknowledgments

The work on natural epivariation was made possible thanks to the genome, methylome, and transcriptome resources provided by the Arabidopsis 1001 Genomes Project. We thank current and past members of the Colot group for discussions and feedback over the lifespan of the project. We thank Edith Heard for critical reading of the manuscript and suggestions for improvement. Computational analyses were performed on the IBENS bioclust computing cluster and we thank the bioinformatics platform team led by Pierre Vincens for their technical support. High-throughput phenotyping benefited from the support of the IJPB Plant Observatory facilities PO-PHENO and PO-VASC.

## Funding

European Union (EpiGeneSys FP7 Network of Excellence #257082, to VC)

French Agence Nationale pour la Recherche (ANR-12-ADAP-00020-01 to VC and OL; ANR-21-CE45-0018, to VC) and by a grant from the PSL-QLife Institute (to VC).

Fondation pour la Recherche Médicale (FRM SPF20170938626 postdoctoral fellowship to PB).

Saclay Plant Sciences-SPS (ANR-17-EUR-0007, to OL).

## Author contributions

Study design: PB, VC

Supervision: PB, VC

Writing manuscript: PB, VC

DNA, RNA, and sRNA extractions of epiRILs as well as ONT-sequencing of 20 strains: EC

Analysis of WGBS-seq data, GWAS and GLM as well as de novo assembly of 20 genomes using ONT-sequencing data: LDO

Epihaplotyping of F9 epiRILs: GBV, VS

Analysis of epivariation segregation over TE-DMRs in the epiRILs and RILs: PB

McrBC analysis of RILs: CX

Analysis of sRNA seq data: AS and PB

Expression and DNA methylation analysis at *HVA22E*: MEM and CX

Single-read DNA methylation analysis using ONT: MBa

Generation of *ddm1* and *ddm1* compound mutants and extraction of sRNAs: FKT and MBo

Estimation of TE family copy-numbers in the genomes of the 1001 Genomes Project: ADF and LQ

Phenotyping of epiRILs using the Phenoscope: EG and OL

## Data and materials availability

Raw BS-seq, RNA-seq, sRNA-seq, and ONT-seq read-data (fastq and fasta of 20 *de novo* genome assemblies, fast5 of epiRIL238) generated and analyzed for this study are available in the European Nucleotide Archive (ENA) under project PRJEB77668.

WGS data was obtained in FASTQ format for the 1047 strains of the 1001genomes.org project from NCBI SRA Project PRJNA273563. Raw RNA-seq data was obtained from NCBI GEO Accession GSE80744. Processed BS-seq data was obtained from NCBI GEO Accession GSE43857. Raw DAP-seq data was obtained from NCBI GEO Accession GSE60143.

## Supplementary Materials

Materials and Methods

Supplementary Text

Figs. S1 to S13

Tables S1 to S11

References (*83–118*)

Datasets S1 to S7

