## Supplementary Materials for "Transposable elements are vectors of recurrent transgenerational epigenetic inheritance in nature"

**The PDF file includes:**

Materials and Methods
Supplementary Text
Figs. S1 to S13
Tables S1 to S11
References

**Other Supplementary Materials for this manuscript include the following:**

Tables S1 to S11
Datasets S1 to S7

### Materials and Methods

Genomic annotations

All genomic analyses were performed using the TAIR10 Col-0 reference genome sequence assembly. The reference TE and gene annotations were used after removal from the latter of the 3,903 TE genes ('transposable_element_gene'; https://www.arabidopsis.org/download/).

Whole-genome Bisulfite sequencing

Single cytosine resolution methylomes were produced using whole-genome Bisulfite sequencing (WGBS-seq, Illumina HiSeq X Ten) of genomic DNA extracted from rosette leaves collected at bolting from pools of six individuals grown in long days, 16h:8h light:dark, at 23°C. The set of 120 *ddm1*-derived epiRILs largely overlaps with that previously epihaplotyped using low-resolution methods(*26*) but were taken here one generation later (F9) and also at generation F21 for 12 of them. Methylome data for the Col-0 WT and fourth generation *ddm1* parental lines were obtained using three biological replicates to provide robust measures of parental methylation levels at individual cytosines. Numbers of sequencing reads reporting methylation or no methylation over each cytosine were determined using Bismark(*83*) and pooled for Cs on opposite strands of CG and CWG (W=A or T) sites, which are symmetrical (in contrast to CCG and CGG sites) and indeed exhibit concordant levels of methylation on the two strands (Fig. S2a).

Identification of Mendelian and reverting differentially methylated sites (DMSs) in the epiRILs

To identify differentially methylated CG and CWG sites (DMSs) we used a generalized linear model (GLM) approach to distinguish in the Bismark output of each site across all three WT and three *ddm1* replicates a significant genotype effect from variation between replicates. To this end, we built for each of the 2.35 millions of CG and CWG sites that are methylated in WT and that have sufficient coverage (≥5 reads per site) in at least 20 epiRILs, a GLM with a negative binomial family and an identity link function using the R package *glm* and the following formula:

MethylatedReads~0+TotalReads+TotalReads:Genotype

where MethylatedReads represents the number of sequencing reads in each sample reporting methylation at the site (pooled between both strands), TotalReads is the sequencing depth at the site (total number of reads supporting both methylation and non-methylation pooled between strands), and Genotype indicates whether the sample is a WT or a *ddm1* replicate. Using this method we identified 807,450 sites with significant p-values for the interaction term (adjusted $p<0.05$ after Benjamini-Hochberg correction for multiple testing) and that are hypomethylated by at least 20% in *ddm1*.

Consistent with previous observations(*22*), most (72.8%) of these DMSs are within annotated TEs or their immediate flanks (300bp on either side, Fig. S1a). Using the same GLM, we then determined the methylation status (i.e. WT-like, *ddm1*-like or intermediate) of these hypomethylated DMSs across epiRILs by comparing the levels calculated in each epiRIL with the predictions from the GLM for each genotype (R function predict.glm): all values below the upper 95% confidence interval (1.96 standard error) for *ddm1* were considered *ddm1*-like, those above the lower 95% confidence interval for WT were considered WT-like, and those that fall between these two thresholds were considered intermediate. For each DMS, we then compared the ratio of WT-like epiRILs across all informative epiRILs to that expected for Mendelian inheritance (73%, given that the F1 individual used to generate the epiRIL population was backcrossed to WT, with 8% of residual selfing(*25*)) . To take into account the number of informative epiRILs, we used a poisson-binomial drawing from the selfing (50-50) or the backcrossing (75-25) distribution: for 20 informative epiRILs the 95% confidence interval of Mendelian segregation is 45%-95% while it narrows down to 64%-82% when the full set of 120 epiRILs is informative (Fig. S1b). Following this approach, we identified 202,721 DMSs (25,1% of total) with WT-like methylation in a percentage of epiRILs consistent with Mendelian inheritance and 587,197 DMSs (72,7% of total) for which this percentage is indicative of either partially or fully penetrant reversion instead. Finally, 17,532 DMSs (2.2% of total) exhibit an excess of *ddm1*-like methylation, which may reflect either *trans* demethylation or else inevitable skewing when the number of informative epiRILs is low (Fig. S1b).

Identification of differentially methylated regions (DMRs)

As expected from previous analyses(*26*), parental DMSs next to each other show coherent epiallelic inheritance across the epiRILs (Fig. S2b), a property that we used to iteratively merge coherently-segregating DMSs into differentially methylated regions (DMRs) as long as they were of the same class (Mendelian or reverting), separated by no more than one DMS of another class, and within 207bp of each other. This threshold distance was chosen as the distance from which the correlation between the methylation status of two adjacent DMSs across epiRILs starts to significantly increase in variability (Fig. S2b). Out of these DMRs we removed those based on less than 5 DMSs and thus obtained 9,422 Mendelian DMRs and 16,040 reverting (Dataset S1).

We then calculated the correlation between the methylation states of each pair of the 7,151 Mendelian DMRs that contain at least 5 DMSs covered in all 120 epiRILs (Dataset S2) to deduce the rate of linkage decay across chromosome arms and pericentromeres (no decay observed in pericentromeres given the scarcity of crossing-overs in these regions). Using a non-linear regression model with variable variance to estimate the expectations of linkage decay as a function of genomic distance - linear in pericentromeres where there is almost no decay (R function nlme::gls) and exponential in chromosome arms (nlme::gnls) - we filtered out 1729 outlier DMRs for which methylation states are either too highly or too lowly correlated (outside of 95% confidence intervals) with other DMRs across the epiRILs (Fig. S2c). High correlation at a distance is observed notably for DMRs overlapping *VANDAL* and *ENSPM* TE annotations, in line with the ability of TEs from these TE families to demethylate cognate TEs in *trans*(*84*) and we therefore removed all the DMRs corresponding to these TE families. Redundant DMRs which are closeby (<5kb) and highly correlated ($R^2$>0.6) were also trimmed down to one representative DMR. For pericentromeric regions, given the high density of DMRs and the scarcity of crossing-overs, we selected between 80 and 100 evenly distributed DMRs for each pericentromere. In total, we obtained a subset of 1,092 evenly distributed and non-redundant Mendelian DMRs (Dataset S3).

Building the epihaplotypic map of 120 F9 epiRILs

To infer the epihaplotypes from the 1,092 DMRs thus defined, we denoised the data using a continuous-time Hidden Markov Model (HMM, R package MSM), with methylation status of DMRs as observed states and the real parental haplotype as hidden states. The initial state occupancy probabilities are the same as the Mendelian expectation (27% *ddm1* vs 73% WT). Using recombination rates measured in female back-crosses(*85*) we calculated the expected recombination rate in our F9 epiRIL population (assuming complete homozygosity by this generation)(*86*). From these recombination rates we calculated the expected transition matrix (WT-derived to *ddm1*-derived and the opposite).

As pericentromeric regions rarely recombine, we used *ddm1*-like and WT-like pericentromeres to calculate the distribution of methylation states of DMRs within *ddm1*-derived and WT-derived intervals, respectively. We used these distributions as emission probability functions to optimize the given transition matrix to adjust for the lack of recombination events in the pericentromeres. Together with the initial state occupancy probabilities and the transition matrix, we used these emission probability functions as parameters in our HMM to derive epihaplotypic intervals delimited by DMRs. Short intervals (<1.5 Mb) were removed from further analysis as well as the small fraction ( 2% ) of larger intervals with DMRs mainly in the intermediate rather than in the *ddm1*- or WT-like methylation state (see e.g. in Fig. S2e), which may correspond to regions of residual heterozygosity. Finally, we noted five epiRILs (#62, #172, #323, #394 and #506) with a high density of *ddm1*-like DMRs in the wild-type derived pericentromeric region of chromosome 4. Remarkably, *VANDAL* and *ENSPM* are also hypomethylated in these intervals, despite the absence of a *trans*-demethylating copy present in a *ddm1*-derived interval. The most parsimonious explanation is that these correspond to *ddm1*-derived intervals with high level or reversion. These intervals were also removed from further analysis.

Overall, the new epihaplotypes (Dataset S3) are more resolutive and extend further than the initial ones (Fig. 1a, S3). Indeed, despite excluding intervals that are poorly resolved or that could reflect residual heterozygosity, we can nonetheless now infer unambiguously the WT or *ddm1* origin ~93% of the genome sequence of each epiRIL on average, compared to 82% before (Fig. S3).

Analysis of DNA methylation levels at TE-DMRs in the epiRILs

To analyze the inheritance of DNA methylation variation at TEs in the epiRILs, we focused on the 7,023 parental DMRs that are ≥100bp-long, comprise at least 10 DMSs (at least 75% of which must be of the same class), overlap annotated TEs, and are located within the newly defined WT and *ddm1*-derived epihaplotypic intervals. Average DNA methylation levels for each C context (CG, CHG, or CHH) within each TE-DMR was calculated within each of the parents and each of the 120 epiRILs as the total proportion of methylated reads summed over all the sites of the given context within the TE-DMR that are covered by at least 5 reads.

To define lower and upper thresholds of WT-like and *ddm1*-like mCG levels, respectively, we took the first percentile of average mCG levels over these TE-DMRs in the three WT replicates and the last percentile in the three *ddm1* replicates (Fig. S1c). The methylation state of a TE-DMR in an epiRIL was considered *ddm1*-like when its mCG level was strictly lower than the former threshold (0.3944), WT-like when it was strictly higher than the latter (0.7586), and intermediate otherwise (Dataset S4). Reversion frequency per TE-DMR was calculated as the proportion of F9 epiRILs that inherited the TE-DMR from the *ddm1* parent and present WT-like mCG levels (counted as 1 event of reversion) or intermediate mCG levels (counted as 0.5 event of reversion). The thresholds between r-low, r-mid, and r-high were set arbitrarily at reversion frequencies of 33 and 66%, respectively. “Stray” TE-DMRs were identified, for non-*VANDAL* and non-*ENSPM* TE families with 4 or more TE-DMRs, as those that remain *ddm1*-like in epiRILs where 75% or more of the TE-DMRs from the same TE family have fully reverted.

Small RNA mapping and analysis

Total RNA was extracted with Trizol according to manufacturer's instructions (Invitrogen) from pools of seedlings of *ddm1* and *ddm1rdr2*, *ddm1rdr6*, and *ddm1dcl2dcl3* compound mutants as well as from inflorescences collected from pools of 5-10 siblings of 10 F9 epiRILs selected for having contrasted epihaplotypic combinations (Fig. 1a, Dataset S3). sRNAs were sequenced by Fasteris with Illumina for the mutants and by the BGI with DNBseq for the 10 F9 epiRILs.

Reads were mapped using bowtie2*(87)* using the --very-sensitive mode (-L 15 -N 1 -p 4) from which unmapped reads and secondary alignments were removed using samtools view (-F 260)*(88)*. The total number of reads ranging in length from 15 to 30nt and mapping on nuclear chromosomes was extracted using samtools view to obtain the sRNA library size of each sample. Multi-mapping reads (attributed randomly to any of its mapping locations by bowtie2 in the default mode) were distinguished from unique-mapping reads based on the presence of an XS:i field in the BAM output of bowtie2 and no-mismatch reads were selected based on the XM:i field. For each category of mapping reads (multi- or single-mapping) the coverage over TE-DMRs in reads of a given length (19 to 26nt-long) was normalized by the sRNA library size of each sample in millions of reads (reads per million, RPM). TE-DMRs were considered to lose all or almost all of their single-mapping no-mismatch 23-24nt sRNAs in *ddm1* when read counts were >0 RPM in WT and in *ddm1* either 0 or at least 99% lower (Fig. S4c). For representation purposes a pseudo-count of 0.001 was added to all read-counts.

To identify all potential sources of multi-mapping sRNAs, multi-mapping reads were aligned, either allowing mismatches or not, at up to 50 locations using the --very-sensitive mode of bowtie2 (-L 15 -N 1 -p 4 -k 50). 23 or 24nt-long reads mapping with or without mismatches over each TE-DMR were then identified using BedTools intersect and their (up to 50) no-mismatch alignments were counted across TE or gene annotations using htseq-count in the -m intersection-strict mode.

QTL$^{epi}$ mapping of reversion at the TE family level

Out of 1092 DMRs used for epihaplotyping, we selected as genetic markers for interval mapping a subset of 139 that are non-redundant, have no missing information, and are separated by >250kb when located within chromosome arms and by >1Mb when within pericentromeric regions (within 5Mb of centromeres) to take into account the lower recombination rates in these regions. These 139 markers (Dataset S3), which cover ~84.4% of the *A. thaliana* genome and are on average separated by 749.7kb, were then used as input for classical interval mapping as implemented in the mqmscan function in R/qtl. For each of the 105 TE families with at least 10 non-systematically reverting TE-DMRs (all but r-0% and r-100%), we used as a trait the average CG methylation level in each epiRIL of the subset of those TE-DMRs that are *ddm1*-derived. The mapping was performed using a step size of 2, and significance was determined for each TE family using 1000 permutations of the data, with LOD significance thresholds corresponding to 5% genome-wide false positive rates. For representation purposes, LOD scores were then normalized by the significance threshold (Fig. S5d). The marker with the highest LOD score was used as top $^{epi}$QTL and the second marker with the highest LOD score that is not within a continuous interval of LOD scores above the significance threshold minus 1 with the top marker was used as second $^{epi}$QTL.

The percentage of variance explained by the epiallelic segregation at each of the $^{epi}$QTLs was estimated by adding iteratively their epihaplotypes as parameters in a logistic GLM. We also used the epihaplotype at the top $^{epi}$QTL in a logistic GLM of average CG methylation levels over the same TE-DMRs but within WT-derived intervals to identify the 24 TE families where it is significantly associated with the epihaplotypic segregation at this locus (p-value<0.05) and thus where effects within *ddm1*-derived intervals may be the result of *trans*-demethylating effects (Fig. S5e).

MRSE-based identification of TE-DMR revertants in large siblings panels

To determine the methylation states of different categories of TE-DMRs in individual siblings of given F9 epiRILs, we obtained from NEB two methylation-sensitive restriction enzymes (MSREs): MspI and AvaII, whose digestion is blocked by DNA methylation and whose restriction sites are typically found once or twice per TE-DMR (MspI | NEB, AvaII | NEB). We then selected TE-DMRs that represented a diversity of non-systematic reversion categories (r-low to r-high) and i) for which we could design primers (Table S2) that specifically amplify a 100 to 200nt-long region containing a digest site of MspI and/or AvaII, ii) were present in contrasted methylation states in the two epiRILs chosen (#92 and #216) and iii) for which we could confirm their differential methylation between 4 Col-WT and 4 Col-*ddm1* plants by comparing digested and non-digested samples using qPCR (Roche LightCycler 480). We then measured methylation states using this method on DNA extracted using Macherey-Nagel 96-well plate extraction kit (Macherey-Nagel, NucleoSpin 96 Plant II) from rosette leaf tissue from 96 F9 2-weeks old siblings of the two epiRILs. All qPCR measures were done in duplicates and calculations were performed on the mean of the two duplicates unless inconsistent (standard deviation >1 Ct), in which case they were discarded. Expected numbers of reversion events were calculated on the basis of the frequency of reversion among *ddm1*-derived F9 epiRILs and assuming a reversion rate per generation that is constant over generations and equal across lines (frequency of non-reversion in F9 = (1-reversion rate)^8).

Phylogeny and divergence of TE sequences

Divergence between related TE-DMRs was calculated by building a phylogeny of all the TE-DMRs within each TE family using phyml (v3.3.3)*(89)* with parameters -s SPR -d nt -a e -q --no_memory_check after trimming the MAFFT (v7.487)*(90)* alignment of fasta sequences using trimal*(91)* in -phylip3.2. For each TE-DMR the distance to the closest TE-DMR was retrieved from the phylogeny of the corresponding TE family using the DendroPy-4.4.0 python library*(92)*.

Identification of spontaneous hypomethylation events in the epiRILs

In a first step, we identified in the set of 120 F9 epiRILs all the cases where a TE-DMR, that do not belong to any of the *VANDAL* or *ENSPM* TE families, present one or two cases of *ddm1*-like hypomethylation within WT-derived epihaplotypic intervals. In order to avoid potentially confounding spontaneous hypomethylation with epihaplotyping errors at the edge of epihaplotypic interval, we then filtered out the 14 such TE-DMRs where hypomethylation is observed at a TE-DMR that is separated from the edge of their WT-derived interval by less than two TE-DMRs that are in a WT-like methylation state.

Epihaplotyping of the 169 epiRILs that minimally overlaps with ours (37 epiRILs are in common) for which single-cytosine resolution methylome data were obtained independently by others*(40)* was performed following the method described by the authors: i.e. based on the WT-like (>0.5) or *ddm1*-like (<0.5) mCG levels of 140 of the 144 DMRs they identified as stably segregating (4 were removed as they deviate by more than 7% from the 75% of WT-like expected). Using this epihaplotypic map, we identified 393 TE-DMRs with one or two events of *ddm1*-like mCG levels within WT-derived intervals that do not belong to the *VANDAL* and *ENSPM* superfamilies. However, we found events of hypomethylation within WT-derived intervals to occur almost systematically on the same epiRILs (pearson correlation > 0.8) for two blocks of 55 and 196 TE-DMRs on Chr3 and Chr5, respectively (Fig. S6e), suggestive of an epihaplotyping error over the corresponding intervals that are most likely *ddm1*-derived. 129 TE-DMRs remained after filtering out the 264 TE-DMRs with pearson correlation>0.8.

Epivariation over TE-DMRs in natural strains

Processed bisulfite sequencing data (BS-seq) and paired-end short reads whole-genome sequencing (WGS) data from leaves of 720 genomes of *A.thaliana* were obtained from the 1,001 Genomes Project (1001genomes.org and signal.salk.edu/1001.php). For each strain, we calculated the average CG methylation weighted by the sequencing depth at each site over TE-DMRs within TEs that are present in the strain based on previous work(*42*). In addition, WGS coverage was calculated using the tool bam-readcount(*93*) in order to detect potential structural variation unaccounted for by sole TE presence/absence polymorphisms. Specifically, for each TE-DMR in each genome, we filtered out regions that have not enough WG-seq coverage (>5% of TE-DMR with 0 coverage) or an excess of reads (median TE-DMR coverage > 2x genome wide median coverage). In addition, we filtered out TE-DMRs with a number of CG sites overly divergent from the reference genome (>50%) or with less than 5 BS-seq reads per CG on average.

Analysis of DNA methylation over TE-DMRs in 20 *de novo* assembled genomes

Long-read Oxford Nanopore Technology (ONT) sequencing data were produced for 20 strains (chosen to represent each genetic group; Table S6) out of the 720 for which BS-seq data is available. High-molecular-weight genomic DNA was extracted from grinded leaves(*94*) and sequenced at ~30X (N50=37kb, Table S6) following recommended guidelines on ONT MinION flow cell (R9.4.1).

Basecalling was performed using Guppy v6.4.6 with the dna_r9.4.1_450bps_sup_plant.cfg model. Long-read fastq files were used for *de novo* assembly using two different assembly tools: Flye(*95*) and NextDenovo(*96*). The assembly with the best N50 was selected. To fix base errors in the assemblies, we polished the genomes using medaka with long-read data (https://github.com/nanoporetech/medaka), and two rounds of NextPolish(*97*) with short-read data. Finally, the contigs of assemblies are scaffolded using Ragtag(*98*) based on alignment on the reference genome. The location of reference genes and TEs were transposed on each genome using Liftoff(*99*), then TE-DMRs were located within their TE by blast. TE-DMRs were considered to be located at a syntenic region in the *de novo* assembly when surrounded by the same reference genes in the genome assembly as in TAIR10. For this analysis we required that at least one the three nearest genes (restricted to TAIR10 IDs ending in 0 to avoid mis-annotated TE-genes that are themselves likely to be translocated between strains) on each side of the TE-DMRs in the *de novo* assemblies were also found surrounding the TE-DMRs in the reference genome. When located in synteny, TE-DMRs were considered full-length when their length in the *de novo* assembly was greater than 80% of that in TAIR10. BSseq data was then remapped on each *de novo* assembly using Bismark(*83*) to calculate mCG levels over the TE-DMRs.

Kinship, local haplotyping, and GWAS

Genome wide IBS and aBN kinship matrix were calculated using the emmax-kin-intel64 function of EMMAX (https://genome.sph.umich.edu/wiki/EMMAX) with parameters -v -s -d 10 and -v -d 10, respectively, on the PLINK-transposed (v1.90p)(*100*) vcf of the short variants identified in the 1001 Genomes (1001genomes_snp-short-indel_with_tair10_only_ACGTN.vcf from https://1001genomes.org/data/GMI-MPI/releases/v3.1/) out of which we only kept bi-allelic SNPs with a minimum MAF of 0.05 and a minimum missing genotyping rate of 0.1 and found within the 720 strains with BS-seq data (SNP720.MAF005.g01 vcf).

Local haplotyping of each TE-DMRs was calculated using BEAGLE(*101*) on the SNP720.MAF005.g01 vcf in windows of 1 kb (500 bp on each side of the most central SNP within each TE-DMR). Local PCAs (Fig. S7l-m) were performed on the SNPs from SNP720.MAF005.g01 located within 1 kb around the TE-DMR.

GWASs were performed using emmax-intel64 with parameters -v -d 10 using the aBN kinship matrix and the SNP720.MAF005.g01 vcf. For epivariation prevalence at each TE-DMR, we only analyzed the 490 naturally epivariable TE-DMRs with at least 5 *ddm1*-like epivariants and with low genomic inflation ($\lambda_{GC}$<1.1). For heatmap representation of the results (Fig. S8a) we subsampled to the SNPs with the highest p-values within 20 kb windows across the genome. For genome-wide epivariation prevalence, we used as phenotype the proportion of natural *ddm1*-like epivariant per strain among all naturally epivariable TE-DMRs present in the strain.

McrBC-qPCR analysis of TE-DMR epivariation in Cvi-0 x Col-0 RILs

We selected for an extended segregation analysis 40 of the 115 naturally hypomethylated TE-DMRs identified in Cvi-0 to represent a variety of superfamilies and reversion categories (Fig. 4c). Genomic DNA was extracted from frozen leaf tissue using Qiagen DNeasy Plant Mini Kit, and 30 to 100ng was used for McrBC (NEB) overnight digestion (15hr). Levels of unmethylated DNA over a target region were calculated as the fraction of DNA concentration measured by qPCR remaining following McrBC digestion compared to the same sample where McrBC is replaced by water (undigested). Efficiency of digestion was confirmed using the *ddm1*-independent DNA methylation at the 3' end of *AT5G13440* as a positive control, and the unmethylated *AT5G13440* as a negative control (primers listed in Table S8).

Using primers designed with primer3 (https://github.com/primer3-org/primer3) to specifically amplify each region (Table S8), we first confirmed for 34 out of the 38 TE-DMRs that they are differentially methylated between Col-0 and Cvi-0 in freshly harvested tissue of two sibling plants. The remaining 4 TE-DMRs either displayed WT-like methylation in Cvi-0 (1 TE-DMR), hypomethylation in Col-0 (1 TE-DMR), or did not amplify in Cvi-0 (2 TE-DMRs), and we therefore excluded them from further analysis. Using McrBC-qPCR, we then measured the methylation levels of the 34 differentially methylated TE-DMRs across a diverse set of 36 F8 Cvi x Col RILs (Table S9), that include the 20 of the minimal set identified by Simon et al. (*47*) as well as 16 additional with maximal genetic differences to the initial set of 20. TE-DMRs were considered hypomethylated when McrBC-qPCR methylation levels were below 40%, WT-like methylated when above 60%, and intermediate otherwise. TE-DMRs and ELF8 alleles were considered as Cvi-derived when both bordering markers are of the Cvi-genotype (B), Col-derived when both are A, and NA otherwise (C or D genotypes or non-agreeing bordering markers). Reversion events were identified as Cvi-derived TE-DMRs with WT-like methylation levels.

Genetic associations between TE-DMR methylation levels and the genetic maps of these RILs(47) were computed using the mqmscan function of the qtl R package with a window.size of 10 and a step.size of 2. The significance threshold at alpha = 0.05 was calculated using mqmpermutation with the same parameters as well as n.perm = 10 and batchsize = 25. LOD scores were normalized by the significance threshold for heatmap visualization (Fig. 4c).

For the 10 RILs for which BS-seq data was publicly available(102), mCG levels were calculated over all naturally epivariable TE-DMRs with *ddm1*-like methylation in Cvi-0 (N=104, Fig. S8e, S10k) or with WT-like methylation in Cvi-0 (N=465, Fig. S10j). Average mCG levels were calculated over both replicates of each RIL and compared to each replicates of the Col-0 and Cvi-0 parental lineages sequenced as part of the same study (Fig. S8e, S10j-k). Hypomethylation and WT-like methylation were classified using the thresholds defined in the epiRILs (<0.3944 for *ddm1*-like and >0.7586 for WT-like). Genotypes were again determined using concordant bordering markers, with reversion events identified as Cvi-derived TE-DMRs with WT-like methylation levels, while spontaneous hypomethylation events as Col-derived TE-DMRs with *ddm1*-like methylation levels. Correlations between methylation levels and genotypes were calculated as a Pearson correlation coefficient.

Generalized linear modeling of genetic determinants

Association between the presence of *ddm1*-like epivariants over the 1068 naturally epivariable TE-DMRs and a set of selected genetic parameters was calculated using a binomial Generalized linear model (logit link) using R glm function. In order to limit the risk of overfitting, we restricted our analysis to the following set of the genetic parameters which we deemed most biologically relevant: copy number of each TE family in each genome, genotype at *ELF8* (using the top SNP at 2:2431793 identified in our GWAS), genotype at *NRPE1* (using the top SNP at 2:16719082 identified in previous studies(42, 52)), presence of an EPICAT as defined previously(36) within, under 500bp, or >500bp from the TE-DMR, distance to the nearest gene, density of CG sites, and length of the TE-DMR. TE family copy-numbers were estimated using average read-depth after remapping short-read sequencing to a library of consensus TEs as described previously(44). For all 10 parameters, the variables were tested as fixed effects in the glm, and for *NRPE1* and *ELF8* genotypes we also included interaction effects with copy-number and gene distance respectively. Discrete parameters such as genotypes or associations with EPICATs were considered as categorical variables, while quantitative parameters were Z-scored (R scale function) to avoid range effects. For each fixed or interaction effect, an individual GLM was built to assess their significance in explaining the prevalence of *ddm1*-like epivariation over the 1068 naturally epivariable TE-DMRs after including the three first components of the PCA of the aBN kinship matrix calculated from the SNP720.MAF005.g01 vcf to take into account population structure (together the three PCs explain ~55% of the variance). If significant ($p<0.05$), the variable with the largest $R^2$ (McFadden adjusted) was included in the GLM, to then iteratively test the effect of the remaining variables. For significant interaction effects, the variable was included along with its fixed effect.

Environmental associations with AraClim biovariables

196 quantitative AraClim biovariables were downloaded from https://gramene.org/CLIMtools/arabidopsis_v2.0/AraCLIM-V2/ and Z-scored (R scale function). To detect potential associations between these biovariables and the occurrence of epivariation at a TE-DMR of interest (among the 490 TE-DMRs with sufficient epivariant counts, see GWAS

methods), we tested each biovariable individually as an additional explanatory variable in a binomial GLM (logit link) which included the first three PCs of the kinship matrix and the two main *trans* modifiers of the number of *ddm1*-like epivariation per strain, namely TE copy-number and *ELF8*. P-values were then corrected using the Benjamini and Hochberg correction for multiple testing using the R p.adjust function. When several AraClim variables were found to be significant (adjusted $p<0.05$), the one explaining the largest fraction of variance (McFadden adjusted $R^2$) was selected.

RNA-seq analyses in natural strains and in the epiRILs

For analysis of gene expression in natural strains, RNA-seq data of the 728 strains analyzed previously*(29)* was downloaded from GEO (Accession GSE80744). For analysis of gene expression in the epiRILs, total RNA was extracted with Trizol according to manufacturer's instructions (Invitrogen) from rosette leaves collected from pools of 5-10 siblings of the same set of 10 F9 epiRILs selected for sRNA analysis and sequenced by the BGI. Reads were mapped using STAR*(103)* with options --outFilterMultimapNmax 1 --alignIntronMax 10000. Coverage over exons was calculated using bedtools coverage and read counts were then normalized over entire transcripts using R package DEseq2. A pseudo count of 0.001 was added to all normalized expression levels to compute log ratios.

For each naturally epivariable TE-DMRs, expression levels of the two closest genes were compared between strains carrying the TE-DMR with *ddm1*-like or with WT-like methylation by computing the log10 ratio between the median expression in the first set over that in the second set of strains (hypo/hyper log ratios) as well as the p-value of a Wilcoxon rank-sum test between the two sets. To obtain random expectations of hypo/hyper log ratios (gray lines in Fig. 6a and S11a) we shuffled randomly the strain labels among all strains with RNA-seq data and repeated the reshuffling for a total of 10 times. For each of the two closest genes, the percentage of variance in expression level that is explained by natural epivariation at the TE-DMR was calculated as the fraction of sum of squares between hypo and hyper strains over the total sum of squares.

For TE-DMRs where epivariation in nature is associated with ≥2-fold expression changes at one of the two nearest genes, we searched in the set of 10 epiRILs sequenced for those that were stably inherited hypomethylated from the *ddm1* parent in at least one of the epiRILs. To limit the possibility of indirect epihaplotypic effects that could be due to the co-occurrence of multiple hypomethylated TE-DMRs in the epiRILs, we restricted our analysis to naturally epivariable TE-DMRs that are within 5kb of the nearest gene and which are not separated from it by another naturally epivariable TE-DMR. For the 99 TE-DMRs matching these criteria we then calculated hypo/hyper log10 ratios of gene expression averages between the epiRILs where they are *ddm1*-derived and hypomethylated and the epiRILs where they are WT-derived and WT-like methylated. Gene expression changes were considered to be congruent between epiRILs and nature (Fig. S11c) if both log ratios were of the same sign and the fold-change was also ≥2-fold in the epiRILs or the p-value in the epiRILs was $<=0.05$. For the TE-DMRs that have reverted to WT-like methylation in at least one but not all of the epiRILs where it is *ddm1*-derived, we also calculated the log10 ratio of gene expression averages between the epiRILs where they are *ddm1*-derived and hypomethylated and the epiRILs where they are *ddm1*-derived and WT-like methylated (rev/hypo log ratios, Fig. S11d).

Phenotypic associations with AraPheno

417 quantitative phenotypic variables were downloaded from the AraPheno database (https://arapheno.1001genomes.org/) and Z-scored (R scale function). For each of them we retrieved from the SNP720.MAF005.g01 vcf the genotypes in the 720 strains with BS-seq data at up to 5 of the top significant SNPs identified in araGWAS (https://aragwas.1001genomes.org/#/). To account first for population structure and the effect of these top SNPs, we built for each Z-scored AraPheno variable a gaussian GLM (identity link) with R glm function using as explanatory variables the first three principal components of the kinship matrix as well as the genotypes at the top significant SNPs (if still significant in the subset of 720 strains). To detect residual associations between the occurrence of epivariation at a TE-DMR of interest (among the 490 TE-DMRs with sufficient epivariant counts, see GWAS methods) and variation at one of these AraPheno, we tested the epivariation as an additional categorical predictor in each GLM. P values were then corrected using the Benjamini and Hochberg correction for false discovery rate using the R p.adjust function and significant associations (adjusted $p < 0.05$) were ranked by the fraction of phenotypic variance explained (McFadden adjusted $R^2$).

*ELF8* sequence analysis

RNA extractions were performed on flash-frozen ground tissue (shoots) from Col-0 WT using Macherey-Nagel Nucleospin RNA plant kit following manufacturer's instructions. Reverse-transcriptase was performed using SuperScript IV (ThermoFisher Scientific) with oligo $d(T)_{20}$ primers and cDNA was amplified by PCR using primers listed in Table S11. The PCR product was gel extracted using Macherey-Nagel Nucleospin Gel and PCR Cleanup kit and sequenced from both primers by Sanger sequencing by Eurofins Genomics (https://eurofinsgenomics.eu/en/custom-dna-sequencing/eurofins-services/tubeseq-services/). The resulting cDNA sequences and the *ELF8'* of Cvi-0 obtained by blast were analyzed and aligned to the *ELF8* TAIR10 sequence using Geneious 10.2.6 (www.geneious.com).

*HVA22E* expression analysis in epiRILs

F10 offsprings of epiRILs #150 and #229 (*ddm1*-like at *AT5TE74320*) as well as of epiRILs #558 and #193 (WT-like at *AT5TE74320*) were obtained from F9 siblings propagated in bulk in standard growth conditions. F10 progeny of each epiRIL was grown on MS-media in Petri dishes in standard growth conditions (23°C) for 2 weeks. Cold-stress (CS) was performed by transferring 2-weeks old F10 seedlings at 4°C for 24 hours. Shoots of seedlings exposed (CS) or not (CTL) to the cold-stress were simultaneously collected at the end of the 24h cold-stress and flash-frozen either individually for methylation analysis or in pools of 8 seedlings for *HVA22E* expression. Methylation levels of *AT5TE74320* were measured in individual F10 seeldings using McrBC-qPCR on flash-frozen shoot tissue as described above (primers listed in Table S8) confirming that the TE-DMR is stably inherited in all F10 offsprings either in the *ddm1*-like (epiRILs 150 and 229) or WT-like (epiRILs 558 and 193) methylation states independently of the cold-stress (Fig. S13b).

RNA extractions were performed on flash-frozen ground tissue (shoots) from five pools of 8 F10 seedlings using Macherey-Nagel Nucleospin RNA plant kit following manufacturer's instructions. Reverse-transcriptase was performed using SuperScript IV (ThermoFisher Scientific) with oligo $d(T)_{20}$ primers and cDNA was cleaned up for qPCR using Macherey-Nagel Nucleospin PCR Cleanup. Expression levels of *HVA22E* were normalized by the average of three reference genes (*AT3G02065, AT2G41020, AT1G13320*). Primers used for qPCR are listed in Table S11.

Phenotyping of epiRILs growth rates in mild-drought or well-watered conditions

Growth and phenotyping of epiRILs on the Phenoscope (https://phenoscope.versailles.inrae.fr/) were performed as described previously(60). Briefly, using seeds obtained from the Versailles Arabidopsis Stock Center (https://publiclines.versailles.inrae.fr/), six plants per epiRIL were grown in individual pots, three of which were maintained at 60% (control) of the maximum soil water content (SWC) and the other three at 30% (mild-drought) of this maximum. First, seeds were germinated at soil saturation (= 100% of the maximum SWC) and seedlings were set up on the robot (= day 0 on the Phenoscope) at 8 days after sowing (DAS). Control and mild-drought SWC were stably reached and maintained from 12 and 16 DAS, respectively. Instant Relative Expansion Rates (RER instant) were calculated as the relative growth rate of the projected rosette area, integrated over +/-3 days windows. The projected rosette area was extracted by segmentation from daily zenithal images.

Statistical analyses

Unless specified otherwise, p-values are obtained by two-sided Wilcoxon rank sum test and statistical analyses and graphics were performed and obtained in MATLAB.

### Supplementary Text

Supplementary Note 1 - Occurrence of *trans* demethylation in the epiRILs

In the 120 F9 epiRILs, TE-DMRs almost always (97.6% of cases) exhibit WT-like methylation when harbored on WT-derived chromosome intervals, with only few instances of intermediate or *ddm1*-like methylation instead (Fig. 1b). Exceptions mainly involve TE-DMRs belonging to the two DNA transposon superfamilies known to encode *trans* demethylation activities(*104, 105*) (*VANDAL* or *MuDR*-like and *ATENSPM*, Fig. S1d, S2e).

Moreover, these superfamilies also represent the bulk (16) of the 24 TE families for which the *ddm1* epigenotype at the $QTL^{epi}$ is associated with lower methylation of WT-derived TE-DMRs (Fig. S5d-e). In addition, these $QTL^{epi}$ intervals often overlap with the location of previously characterized active *trans*-demethylating elements(*84*) (Fig. S5d). Hence, the $QTL^{epi}$ identified for these superfamilies appear to have a dual effect, favoring either reversion when WT-derived or *trans*-demethylation when *ddm1*-derived and we therefore excluded them from further analysis.

Supplementary Note 2 - Intermediate DNA methylation in the epiRILs reflects incomplete reversion at the single-molecule level

To investigate whether intermediate DNA methylation observed in epiRILs by F9 reflect the presence of a mixture of progeny with or without reversion, or rather incomplete restoration of WT-like levels in individual progeny we performed for one epiRIL chosen at random long-read ONT sequencing and single-molecule analysis of DNA methylation of the same DNA sample used for BS-Seq, which was obtained from a pool of six seedlings (Fig. S1f-g).

In order to enrich the ONT sequencing power over regions of interest, we used the adaptive sampling option available on the GridION sequencer (Oxford Nanopore Technologies). High-molecular-weight genomic DNA (>20 kb-long fragments) was extracted from grinded leaves of the same pool of 6 epiRIL238 F9 plants used for BS-seq following a protocol adapted from *Mayjonade et al, 2016* (94) for bigger extraction volumes. In order to reduce the library size for adaptive sampling, DNA was first sheared at 10 ng.uL-1 by sonication for a single 30s interval at the lowest sonication intensity on a standard Bioruptor (Diagenode) to obtain a range of DNA fragments from 200 bp to >20 kb. Sonication was followed by size selection with Sera-Mag SpeedBeads Carboxylate (GE Healthcare)(*106*), to eliminate fragments smaller than ~1500 bp. Sequencing was performed on a LSK109 flow cell following manufacturer's procedures and using for adaptive sampling a 1.07 Mb-long set of target regions (~1% of reference genome) that overlap by at least 5 kb the 10 TE-DMRs localized in *ddm1*-derived intervals in epiRIL238 with intermediate DNA methylation levels as well as 45 TE-DMRs that have fully reverted (Table S3). mCG levels were calculated over TE-DMRs using DeepSignal-plant(*107*) either at the regional level or at the single-read level over the TE-DMR interval for reads that cover >50% of the TE-DMR.

Out of the 10 TE-DMRs with intermediate methylation in this epiRIL, we found that it reflects for 9 of them progressive remethylation at the level of individual progeny rather than a mixture of non-revertant and fully revertant siblings (Fig. S1f-g). This is in line with previous work indicating that reversion to WT methylation within an epiRIL often occurs in a stepwise manner across several generations(*33*) and with our observation that intermediate methylation at F9 usually translates into WT methylation by F21 (Fig. 1c).

Supplementary Note 3 - EPICATs antagonize reversion

The two closely related TE-DMRs of the *ATGP2N* family located on chromosomes 1 and 4 provide a striking illustration of the negative impact of these EPICATs on reversion (Fig. 3c). Specifically, despite an abundance of multi-mapping 23-24nt sRNAs matching the two TE-DMRs in epiRIL #232, where both derive from *ddm1*, the copy present on chromosome 1 is fully reverted by F9 whereas that on chromosome 4, which contains a strong and long EPICAT, remains hypomethylated. In fact, whether stray or not, TE-DMRs that are associated with EPICATs revert less frequently (Fig. S6c) and appear to require for reversion higher levels of matching multi-mapping 23-24nt sRNAs (Fig. 3d) than the other TE-DMRs of the same family. Thus, inheritance of *ddm1*-induced hypomethylation is significantly favored when there is a strong Pol II transcriptional activity at or near the TE-DMR.

Supplementary Note 4 - *ddm1*-like epivariants in nature are not the result of cryptic structural variation

Given that structural variants are difficult to detect using short read-sequencing and may generate false calls of *ddm1*-like hypomethylation, we produced *de novo* genome assemblies using long-read ONT sequencing for 20 genetically diverse strains (Table S6) to verify that naturally epivariable TE-DMRs match those studied in the epiRILs (see Methods). Using this approach, we determined that on average, 18% of TE-DMRs in any strain are either truncated or missing from the reference location. However, this percentage drops to 13% for the set of TE-DMRs with natural epivariation (Dataset S6). Moreover, whether the WGBS-seq data are mapped onto the *de novo* assemblies or the reference TAIR10 assembly ($R^2$=0.86), methylation calls for the 82% of TE-DMRs with no structural rearrangement are highly congruent (>95%,Fig. 4b). Together, these observations indicate that our stringent filtering step considerably reduced the number of spurious natural epivariants we may have otherwise called because of the presence of related TEs in the non-reference strains and the mismapping issues it can generate(*46*).

Supplementary Note 5 - Case of *trans* association in RILs reflects TE insertion polymorphism between Col-0 and Cvi-0

For one TE-DMR, carried by an *ATCOPIA23 (AT3TE90530)* on chromosome 3 in Col-0, associations are observed in *cis* at the location of the TE-DMR on Chr3 but are strongest with markers in *trans* on Chr5 (Fig. 4e). This *trans* association matches with a TE insertion polymorphism of *ATCOPIA23* we had previously identified in Cvi-0 using short-reads(*42*), which we could readily confirm by aligning the Cvi-0 ONT assembly to the Col-0 genome at this location (Fig. S8d, bottom panel). Moreover, by re-mapping BS-seq reads on the Cvi-0 ONT assembly we found in addition that Chr5 *ATCOPIA23* insertion is in a hypomethylated state (Fig. S8d, bottom panel). In contrast, at its original location on Chr3 where it is fully methylated in Col-0 we found that *ATCOPIA23* is altogether absent in Cvi-0 (Fig. S8d, upper panel). Altogether, this TE insertion polymorphism fully explains the dual association of hypomethylation with markers in *cis* and in *trans*.

Supplementary Note 6 - Functional analysis of *ELF8* alleles

The TAIR10 sequence of *ELF8* contains a frame-shifting single-nucleotide deletion within exon 6 (indicated with a * in Fig. 5e and S10a) which we could not confirm by sequencing ~1kb of the cDNA of the Col-0 *ELF8* allele surrounding this deletion. Instead, we found at this position a single-G insertion, which results in a perfectly in-frame coding-sequence (Fig. S10a). Therefore, we considered the single-nucleotide deletion as a sequencing error in TAIR10 and corrected accordingly the predictions of the Col-0 protein sequence (Fig. S10b). We then used this corrected sequence as $ELF8^{ref}$ for comparisons with that of the *ELF8'* allele carried by Cvi-0 (Fig. 5e, S10b-c).

Although the coding-sequence of the *ELF8'* allele carried by Cvi-0 differs from that of the reference allele by 40 SNPs, none of these are missense (Fig. 5e, S10b). Moreover, the seven SNPs that are non-synonymous do not alter appreciably the predicted folding of the protein sequence. Comparison of the predicted folding of the protein sequence of both alleles was performed using AlphaFold2(*108, 109*) finding no appreciable alteration (Fig. S10c), which suggests therefore that the ELF8' gene product is fully functional.

Further supporting this conclusion, we found that the higher expression levels of *ELF8* in strains carrying *ELF8'* rather than $ELF8^{ref}$ observed in matching published RNAseq data(*29*) (Fig. 5g) are associated with an overexpression of both *FLOWERING LOCUS C* (*FLC*) and *GENERAL REGULATORY FACTOR11* (*GRF11*) in the same set of strains (Fig. S10d-e). In line with the higher expression levels of *FLC, ELF8'* accessions also tend to be later flowering (Fig. S10f) according to published FT16 phenotypic data of an overlapping set of strains(*28*). As a direct role for ELF8 in the establishment and maintenance of H3K4 methylation over promoter regions has been obtained for both *FLC* and *GRF11* (*110–112*) , these observations are strongly indicative of *ELF8'* being an over-expressed functional allele.

Moreover, we observed lower mCHH and mCG methylation levels in strains carrying the *ELF8'* rather than the $ELF8^{ref}$ allele over TEs that lose CHH methylation in the *jmj14 ldl1 ldl2* triple mutant that is defective for the three H3K4 demethylase activities(*48*) (Fig. S10g-h). As a strong reduction of RdDM targeting of TE sequences within ~2kb of H3K4me2/me3-marked genes has been observed in this mutant (*48*), these observations further suggest a role of *ELF8* in antagonizing RdDM near genes.

### Supplementary Note 7 - Role of *ELF8* alleles in RILs

To investigate in more detail the potential causal relationship between genetic variation at *ELF8* and epivariation, we took advantage of the fact that Cvi-0 carries the *ELF8'* allele and asked if the number of reversion events we detected in the 36 Cvi-0 x Col-0 RILs (Fig. 4e) correlates negatively with it. Unexpectedly, 80% of reversion events took place in RILs that harbor the *ELF8'* allele, a significant enrichment compared to the number of RILs that inherited the corresponding TE-DMRs from the Cvi-0 parent (fisher test p=0.01; Fig. 4e). Moreover, we observed a similar bias in the 10 RILs that were analyzed using BS-seq (81% of 27 events, p=0.002) and no reversion in the Cvi-0 parental line itself (Fig. S10j). Although it remains to be determined why *ELF8'* is associated with increased reversion to WT-like DNA methylation in the RILs, this finding effectively rules out a major role of ELF8 activity on its own in stabilizing hypomethylated epialleles. To determine if ELF8 may favor instead the occurrence of spontaneous TE epivariation, we examined the DNA methylation status of all 1,068 naturally epivariable TE-DMRs in the 10 Cvi-0 x Col-0 RILs with methylome data (see Methods). We detected only two spontaneous hypomethylation events, in agreement with the rate of regional loss of DNA methylation over TE sequences in classical mutation accumulation lines derived from fully WT parents(*113*), and neither epivariant was in the *ELF8'* background (Fig. S10j-k). Therefore, we can also exclude a major contribution of ELF8 activity by itself in the generation of spontaneous epivariation. Together, these findings indicate that although natural variation at *ELF8* is an important determinant of the differences in the prevalence of natural epivariants between strains, other factors, either genetic or environmental, likely affect their occurrence or stabilization.

### Supplementary Note 8 - Limitations of the study

Our study likely underestimates the extent of natural *ddm1*-like epivariation at TEs given the stringent filters we used to define it. Moreover, we did not consider TEs not present in the reference Col-0 genome, which typically correspond to recent, large-effect insertions(*42*) and which can also epivary in nature(*114*). Furthermore, *ddm1*-induced TE-DMRs do not include, by design, any of the ~25% of TEs that are already hypo- or un-methylated in WT Col-0(*115*). This limitation effectively biased our GWAS and GLM approaches towards finding genetic modifiers that increase rather than decrease the prevalence of epivariation. Also, while TEs are implicated in most known examples of epiallelic variation with phenotypic consequences in plants, DNA methylation variation affecting other types of sequences, such as non-coding RNA genes and some protein-coding genes, may also contribute to phenotypic diversity in Arabidopsis(*116–118*).

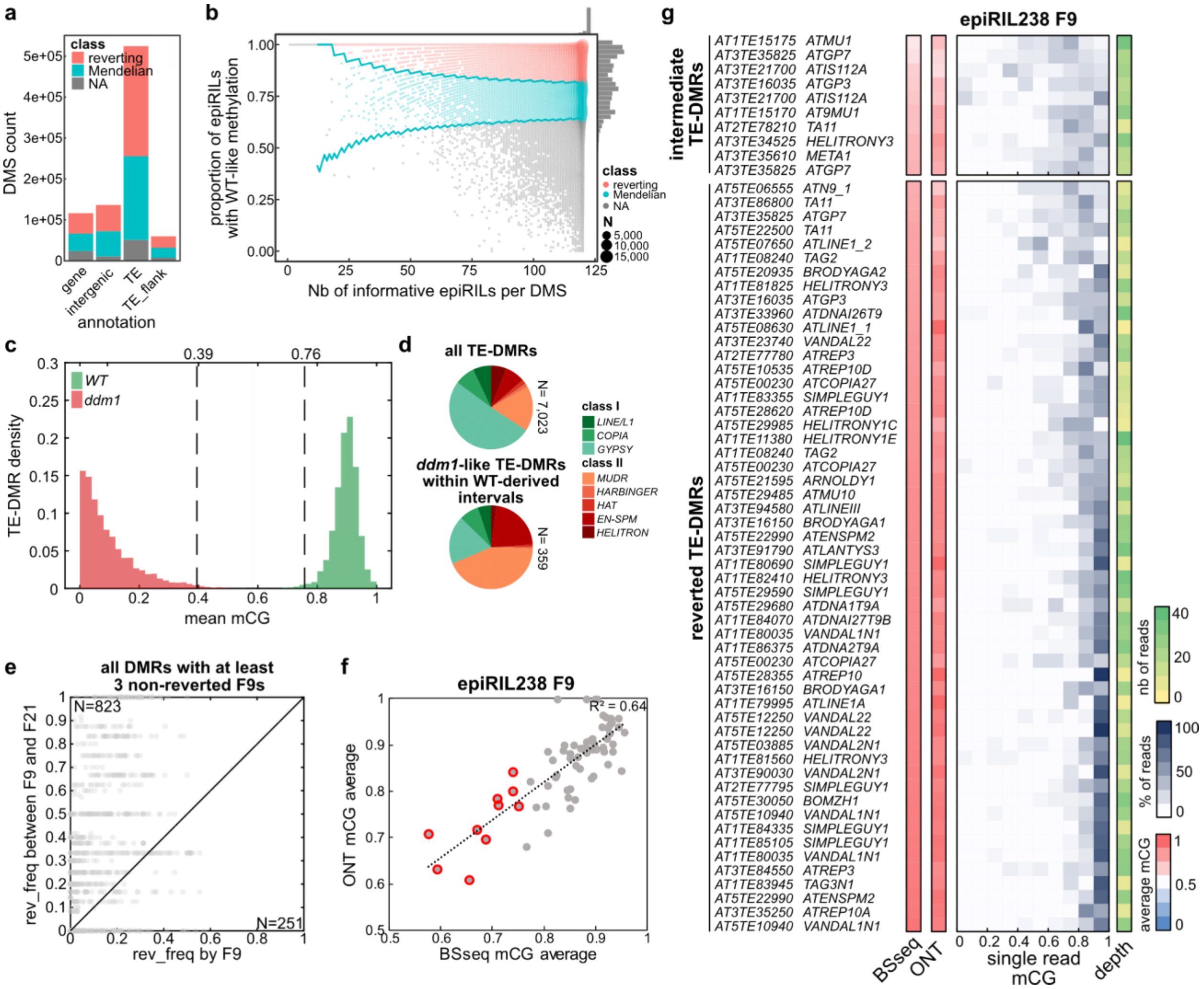


**Figure S1.**

(a) Distribution of Mendelian and reverting DMSs within genic, intergenic, TE, and TE flanks, based on TAIR10 annotation. (b) Proportion of epiRILs with WT-like methylation per DMS as a function of the number of informative epiRILs at each DMS. 95% confidence intervals for Mendelian inheritance using a poisson-binomial are represented as cyan lines within which DMSs are considered as Mendelian, with 8% of selfing estimated from (25) . DMSs with higher proportions of WT-like epiRILs are considered as reverting. Frequency distribution of DMSs in terms of each metric are represented on the corresponding sides. (c) Distribution of mean mCG in *ddm1* (3 replicates) and WT (3 replicates) over the 7,023 TE-DMRs. The values of the last and first percentiles, respectively, are indicated by dashed lines. (d) TE superfamilies contributing to all 7,023 TE-DMRs (left pie chart) or to the TE-DMRs with *ddm1*-like mCG levels within WT-derived intervals in F9 epiRILs (right pie chart). (e) Reversion frequency among F21 epiRILs over TE-DMRs that are non-reverted in at least 3 F9s plotted against reversion frequency among F9 epiRILs. (f) Average mCG levels calculated by BS-seq vs ONT over 10 TE-DMRs with intermediated DNA methylation levels (based on BS-seq) within *ddm1*-derived intervals in epiRIL238 F9 (circled in red) as well as over 45 TE-DMRs that have fully reverted. (g) Distribution of single-read mCG levels calculated by ONT over the same set of TE-DMRs in epiRIL238 F9.

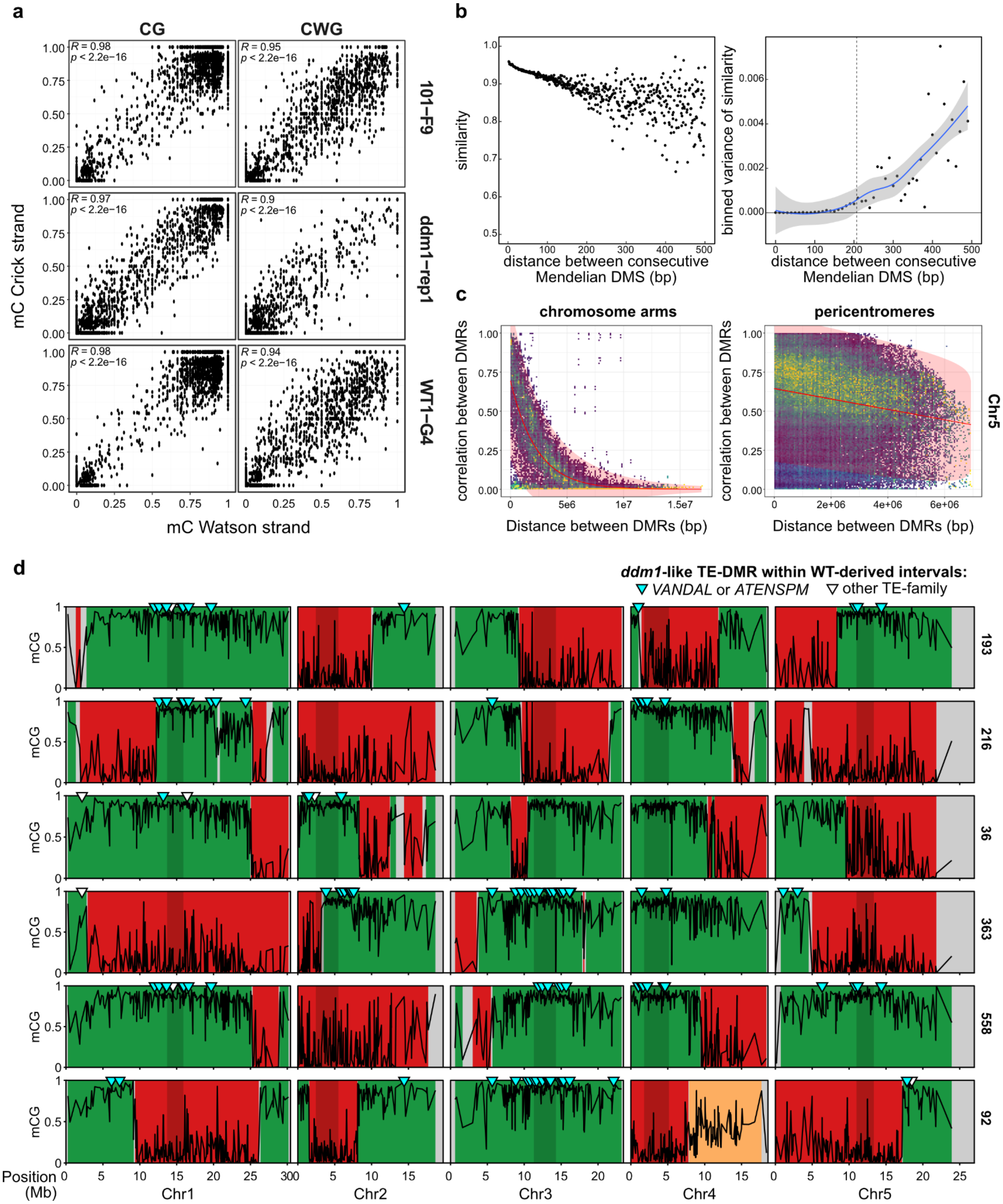


**Figure S2.**

(a) mC levels on Crick vs Watson strands for the CG and CWG contexts in 1 randomly chosen F9 epiRILs, as well as in 1 replicate of the *ddm1* and WT parental lines. (b) Similarity between consecutive Mendelian DMSs as a function of the distance separating them (left panel). Similarity was calculated as 1 - the normalized Manhattan distance, where *ddm1*-like, intermediate and wt-like categories were given integer scores 1, 2 and 3 respectively to calculate a Manhattan distance, which was then divided by 120 (maximum possible distance, signifying perfect disagreement between the methylation states of two DMS across the epiRILs) for normalization. Variance in similarity between consecutive Mendelian DMSs as a function of the distance separating them (right panel). Lowess regression and 95% confidence intervals are represented in a blue line and gray area, respectively. The distance at which variance gets significantly higher than 0 (outside 95% confidence interval) is indicated by a dashed line (207bp). (c) Pearson squared correlation between pairs of DMRs as a function of the distance separating them within chromosome arms and pericentromeres of chromosome 5 (the distance between two markers on either side of a centromere is their distance minus the length of the centromere). Regression and 95% confidence intervals are indicated in red. (d) Mean mCG levels over the 1,092 DMRs (black line) used for the HMM reconstruction of parental epihaplotypes (WT-derived in green, *ddm1*-derived in red, NA in gray, potential residual heterozygosity in orange) in 6 of the 10 F9 epiRILs sequenced for mRNA and sRNA analysis. Events of *ddm1*-like hypomethylation over TE-DMRs within WT-derived intervals are indicated by a cyan arrow if carried by a *VANDAL* or *ATENSPM* TEs or a white arrow otherwise.

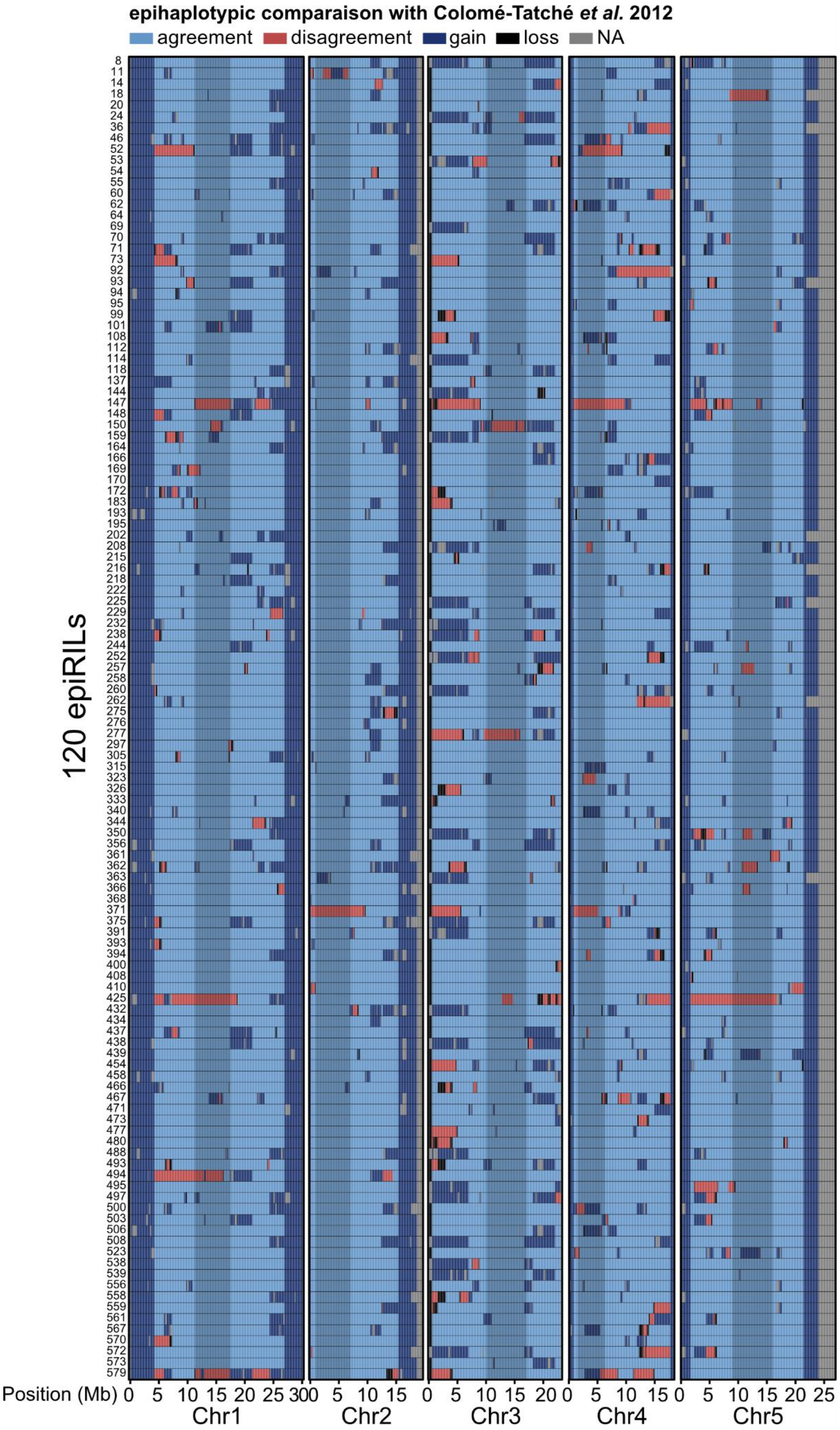


**Figure S3.**
Comparison between epihaplotypes of this study and those obtained in Colomé-Tatché et al. (26) for the same set of 120 epiRILs.

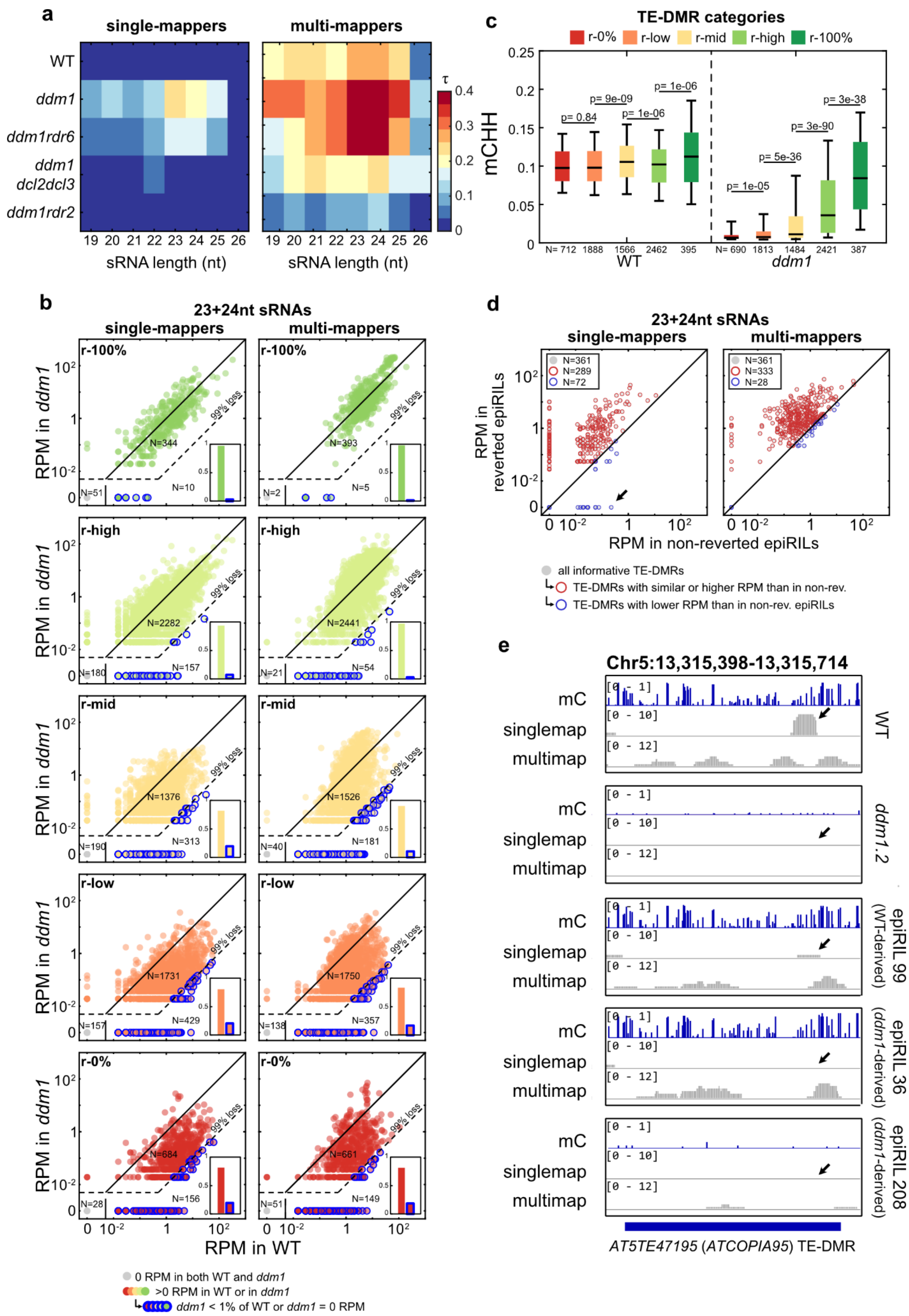


**Figure S4.**

(a) Correlation (Kendall τ) between single-mapping or multi-mapping sRNA levels (RPM) and reversion frequency in WT and different mutant backgrounds. (b) Scatterplot of single-mapping (left) or multi-mapping (right) 23+24nt-long sRNA read counts (RPM) over TE-DMRs of each reversion category (from r-0% to r-100%) in *ddm1* vs in WT. TE-DMRs that lose >99% or all single-mapping sRNA in *ddm1* compared to WT are highlighted in blue and their proportion out of TE-DMRs that have non-zero read-counts in at least one sample (non-gray) are represented as barplots in the bottom right corner. (c) Levels of mCHH in WT and *ddm1* over the five TE-DMR reversion categories. (d) Scatterplot of single-mapping (left) or multi-mapping (right) 23+24nt-long sRNA read counts (RPM) over non-systematically reverting TE-DMRs (r-low, r-high, and r-mid) that lose >99% or all single-mapping sRNA in *ddm1* compared to WT (blue circles in S4b) in epiRILs where they are reverted vs in epiRILs where they are non-reverted. TE-DMRs that have higher read-counts in reverted than in non-reverted epiRILs are highlighted in red while those where it is lower or equal are in blue. The black arrow points to TE-DMRs that revert without regaining single-mapping sRNAs. (e) DNA methylation levels and abundance of matching 23-24nt sRNAs over the *ATCOPIA95* TE-DMR (*AT5TE47195*) in WT, *ddm1*, epiRIL36 (where it was inherited from *ddm1* and had reverted without regaining the single-mapping sRNAs lost in *ddm1),* epiRIL99 (where it was inherited from WT), and epiRIL 208 (where it was inherited from *ddm1* and had not reverted). The black arrow points to single-mapping sRNAs produced in WT.

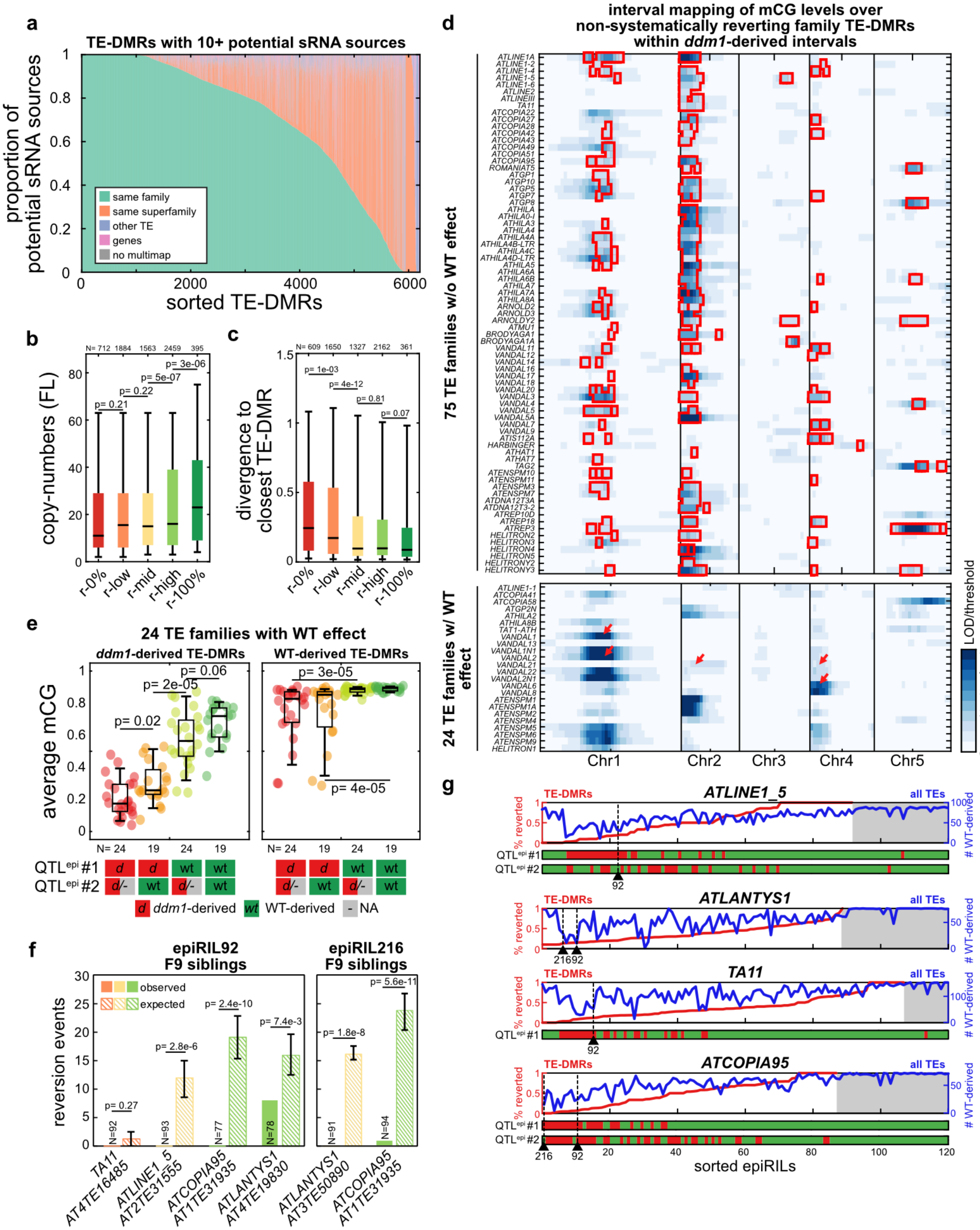


**Figure S5.**

(a) Proportion of potential 23-24nt sRNA sources for each TE-DMR with 10+ potential sources that correspond to TEs of the same family, superfamily, or to other TEs or to genes. (b) Copy-number of full-length (FL) TEs across categories of TE-DMRs. (c) Divergence to closest TE-

DMR across categories of TE-DMRs. (d) Heatmap of the normalized LOD scores from the interval mapping of the average *ddm1*-derived mCG levels by TE family. For the 75 TE-families with a significant $^{epi}$QTL without WT-effect (top panel) the intervals with significant LOD scores encompassing a potential 23-24nt sRNA source are highlighted in red. For the 24TE-families with an $^{epi}$QTL with a significant WT-effect (bottom panel) the location of active *trans*-demethylating elements previously characterized *(84)* is indicated by red arrows. (e) Average mCG levels for each TE family with a WT-effect over *ddm1*-derived or WT-derived TE-DMRs found in epiRILs with the WT or *ddm1* epihaplotypes of QTL$^{epi}$ #1 or #2. (f) Observed numbers of siblings with WT-like DNA methylation (i.e. reversion events) measured by targeted MSRE-qPCR for 4 TE-DMRs in >75 siblings of two F9 epiRILs in which these TE-DMRs are inherited from and *ddm1*-like. Colors follow the reversion categories of 1b-c, with full and striped bars indicating respectively the observed and expected number of reversions. (g) For each TE family of the TE-DMRs analyzed in S5f, fraction of reverted TE-DMRs per epiRIL (red), number of WT-derived TEs per epiRIL (blue), and epihaplotypic origin (*ddm1* in red and WT in green) of potential sRNA sources identified within top QTL$^{epi}$, if identified, across the 120 epiRILs sorted by increasing fraction of reverted TE-DMRs. epiRILs for which no TE-DMR is inherited from *ddm1* are highlighted in gray.

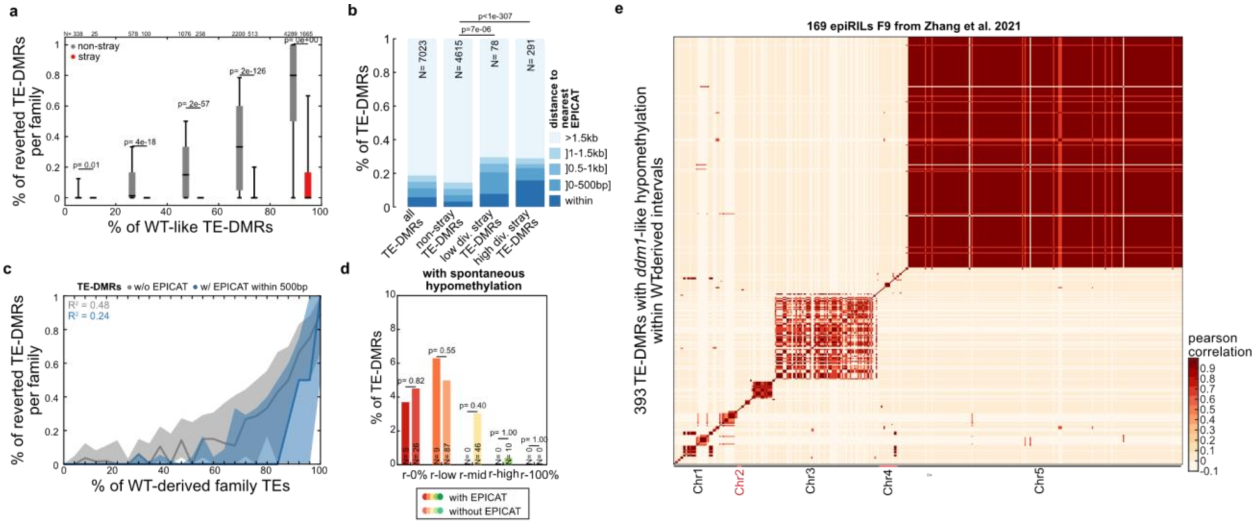


**Figure S6.**
(a) Percentage of reverted TE-DMRs per TE-family, ordered as a function of the percentage of WT-like TE-DMRs for stray and non-stray TE-DMRs. (b) Percentage of TE-DMRs by categories of distance to nearest EPICAT for all, non-stray, "low div." stray, and "high div." stray TE-DMRs. (c) Percentage of reverted TE-DMRs per TE-family as a function of the percentage of WT-derived TEs in the epiRILs and according to whether TE-DMRs are located near (<=500bp) or overlap an EPICAT, or none of these. (d) Proportion of TE-DMRs with spontaneous *ddm1*-like hypomethylation in both epiRIL datasets across r-0% to r-100% reversion categories depending on whether or not they are located near or overlap and EPICAT. The p-values indicate the result of the Fisher exact test between both categories for each reversion category. (e) Heatmap of Pearson correlation in the distribution of the *ddm1*-like hypomethylation events detected within WT-derived intervals the 169 epiRILs from Zhang et al. *(40)*. The two large blocks of strong correlations seen in the right panel suggest epi-haplotyping errors, with all of the concerned TE-DMRs deriving from *ddm1* rather than the WT parental line and were subsequently excluded from analysis.

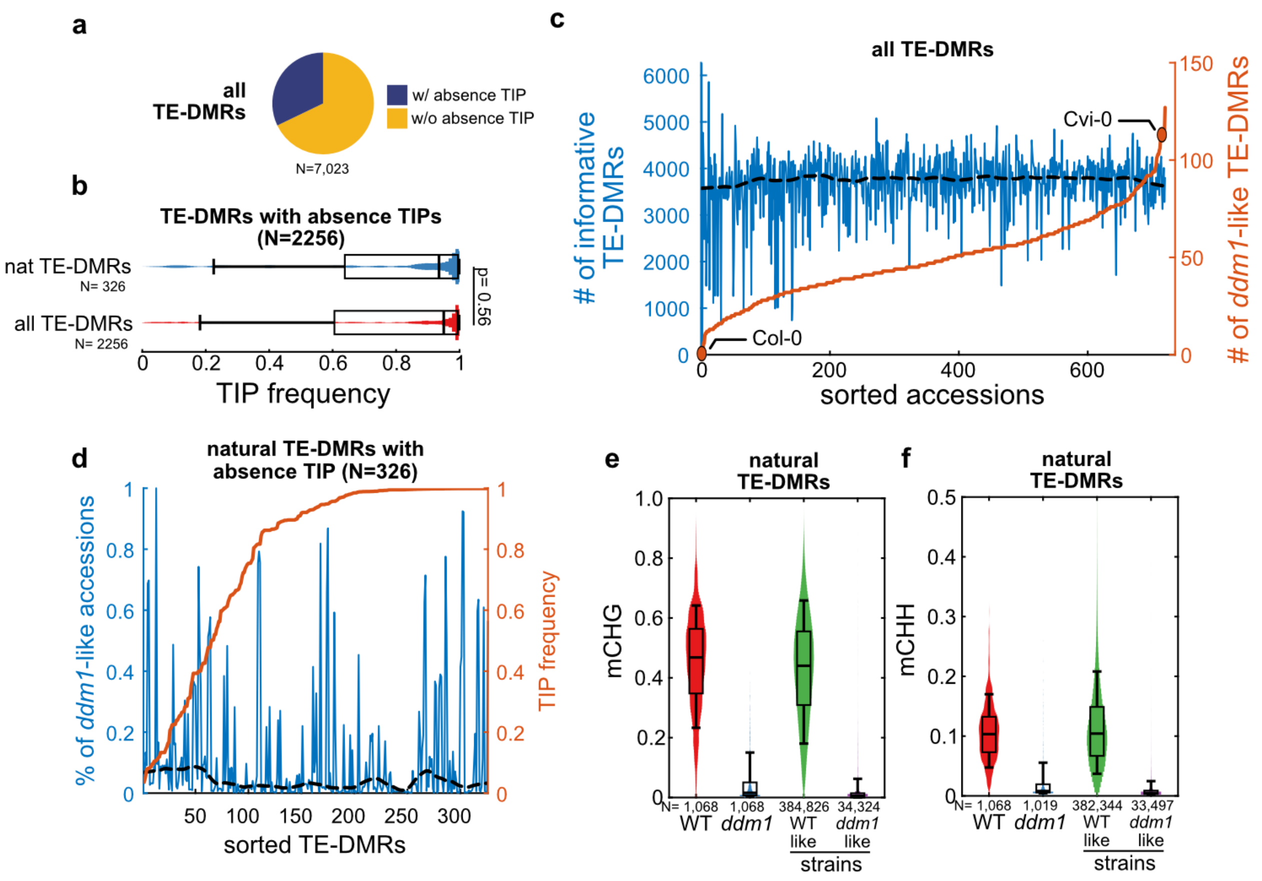


**Figure S7.**
(a) Proportion of TE-DMRs for which an TE insertion polymorphism (absence TIP) was detected among the 1047 genomes analyzed previously(42). (b) Frequency across the 1047 genomes of the TIPs corresponding to all TE-DMRs analyzed or those that are naturally epivariables (nat. TE-DMRs). (c) Number of informative TE-DMRs per genome (blue) and LOESS regression (dotted line) with strains being sorted by the number of epivariants they carry (orange). (d) Proportion of carriers with epivariation per naturally epivariable TE-DMRs with an absence TIP (blue). TE-DMRs are sorted by TIP frequency (orange). The dotted line represents the LOESS regression of the proportion of carriers with epivariation across TE-DMRs. (e,f) Comparison of mCHG and mCHH levels over naturally epivariable TE-DMRs in Col-0 WT and *ddm1* vs in strains without or with epivaration.

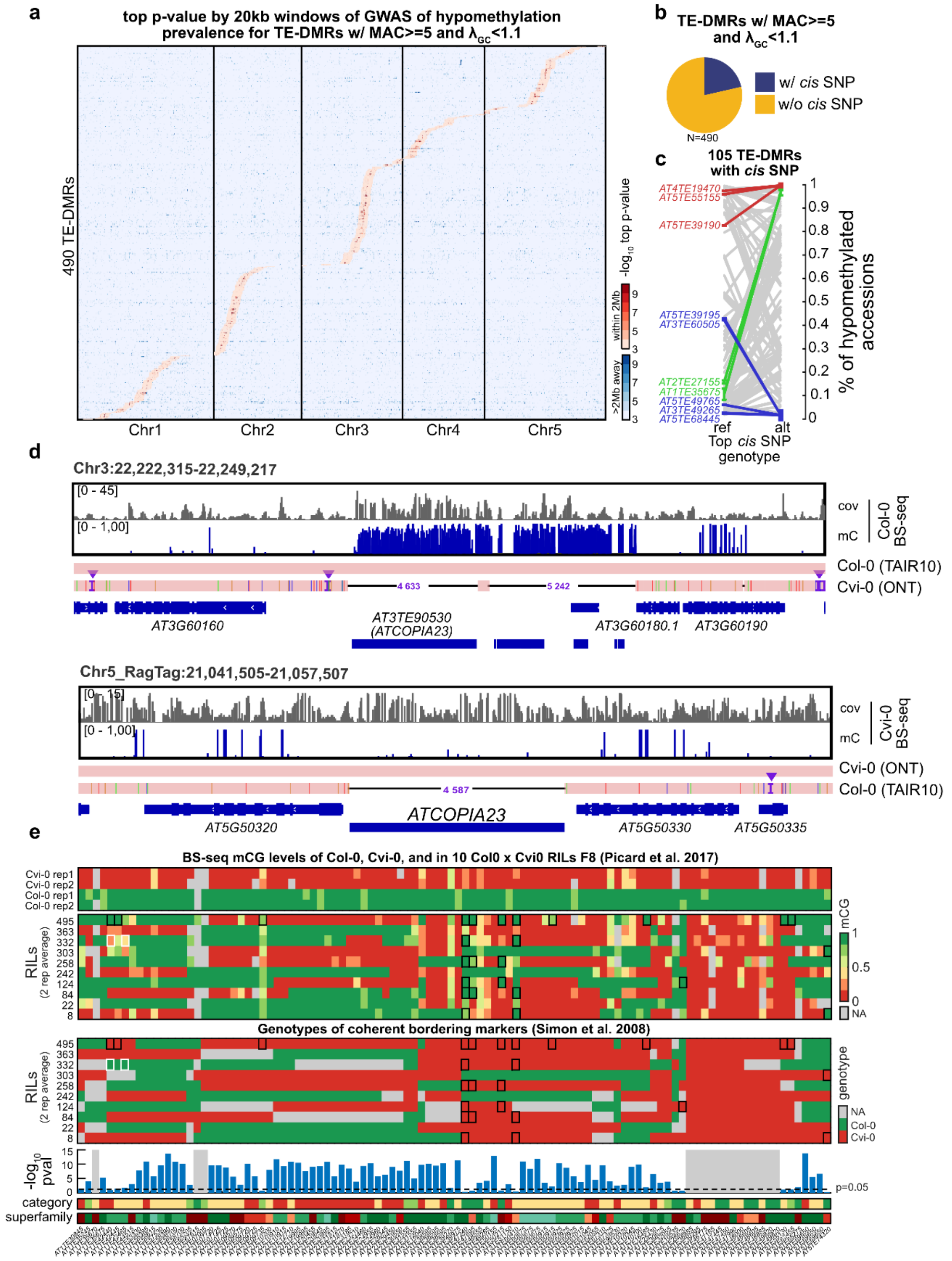


**Figure S8.**

(a) Heatmap of GWAS top p-values by 20kb windows for 490 TE-DMRs with sufficient minor epivariant counts and low genomic inflation ($\lambda_{GC}$<1.1). Windows near TE-DMR are colored in shades of red. (b) Proportion of 490 TE-DMRs with and without significant SNPs found within *cis* 20kb window. (c) Proportion of strains with epivariants that carry the reference or alternate nucleotides at the top *cis* SNP for each of the 105 TE-DMRs with *cis* SNPs. TE-DMRs with no or only accession with epivariation per genotype are highlighted in blue or red, respectively. TE-DMRs with a ≥80% difference between genotypes in the proportion of strains with the epivariant are highlighted in green. Error bars represent the standard-deviation in 10 random subsamples of two third of the strains. (d) Alignment of the Cvi-0 genome assembly against TAIR10 in the region of Chr3 surrounding *AT3TE90530* (*ATCOPIA23*), which is absent from Cvi-0 (upper panel). Alignment of TAIR10 against the Cvi-0 genome assembly in the region of Chr5 surrounding the insertion of *ATCOPIA23* that is present in Cvi-0 but absent from Col-0 (lower panel). For each region the BS-seq coverage and mC levels are represented above the alignments. (e) Heatmap of mCG levels (BS-seq) in 10 Cvi-0 x Col-0 RILs(*102*) of 104 TE-DMRs with epivariation in Cvi-0 in comparison with genotypes at flanking markers (lower panel). Reversions events are highlighted in black squares and spontaneous epivariation in white squares. When calculable, the p-value of the Pearson correlation is represented for each TE-DMR as a bar chart. Reversion categories and TE superfamilies are indicated below for each TE-DMR (legend as in Fig. 1, S1).

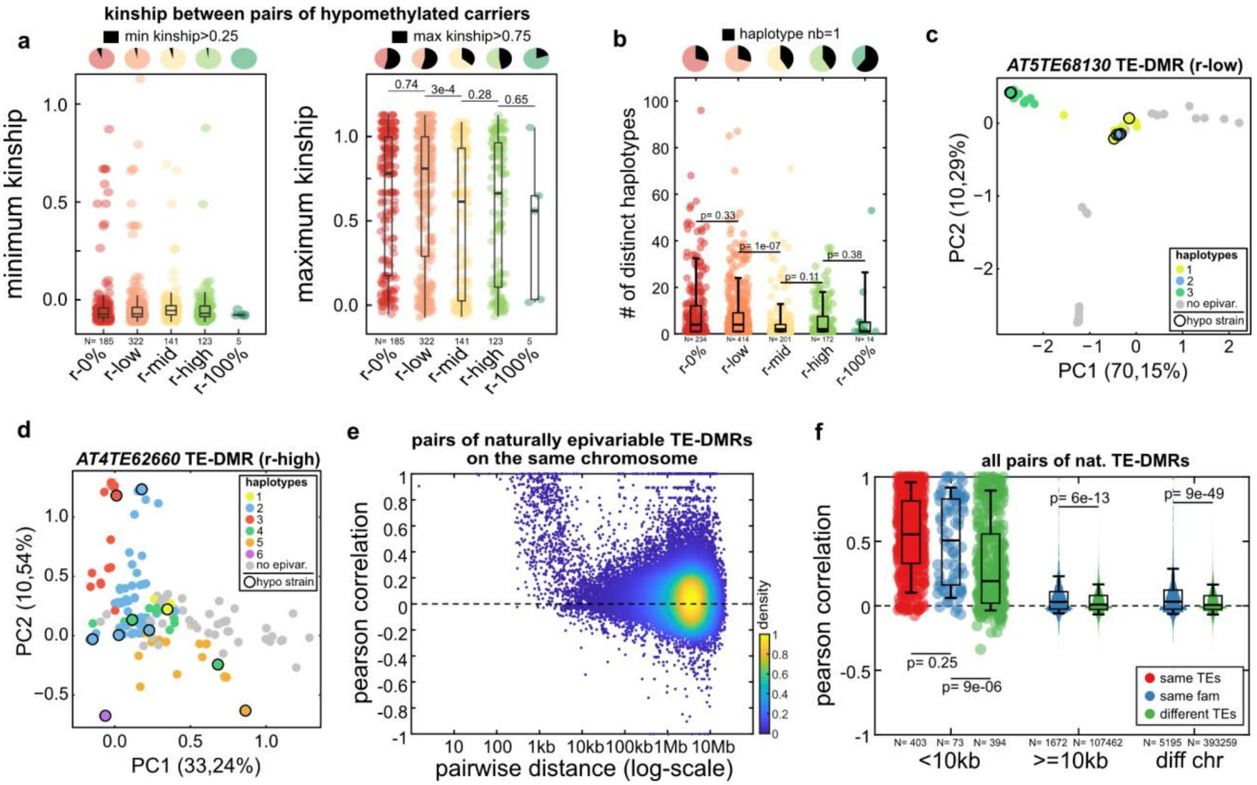


**Figure S9.**

(a) Minimum and maximum kinship observed between any two strains carrying an epivariant of the same TE-DMR, ordered by reversion category. Pie charts indicate for each reversion category the fraction of TE-DMRs that have minimum kinship above 0.25 and maximum kinship above 0.75. (b) For each reversion category, number of distinct haplotypes carrying an epivariant for the same TE-DMR. Pie charts indicate in each case the fraction of TE-DMRs that have only one haplotype per epivariant. (c,d) PCA of SNPs within 1kb windows of representative r-low and r-high TE-DMRs. Strains are colored by haplotype if at least one carrier strain has an epivariant for the TE-DMR considered and are in gray otherwise. (e) Pearson correlation between pairs of naturally epivariable TE-DMRs located on the same chromosome versus the distance between the two TE-DMRs. (f) Pearson correlation between all pairs of naturally epivariable TE-DMRs that are either within 10kb of each other, further away on the same chromosome, or on different chromosomes, whether they are carried by the same TEs, or TEs of the same TE family, or by different TEs.

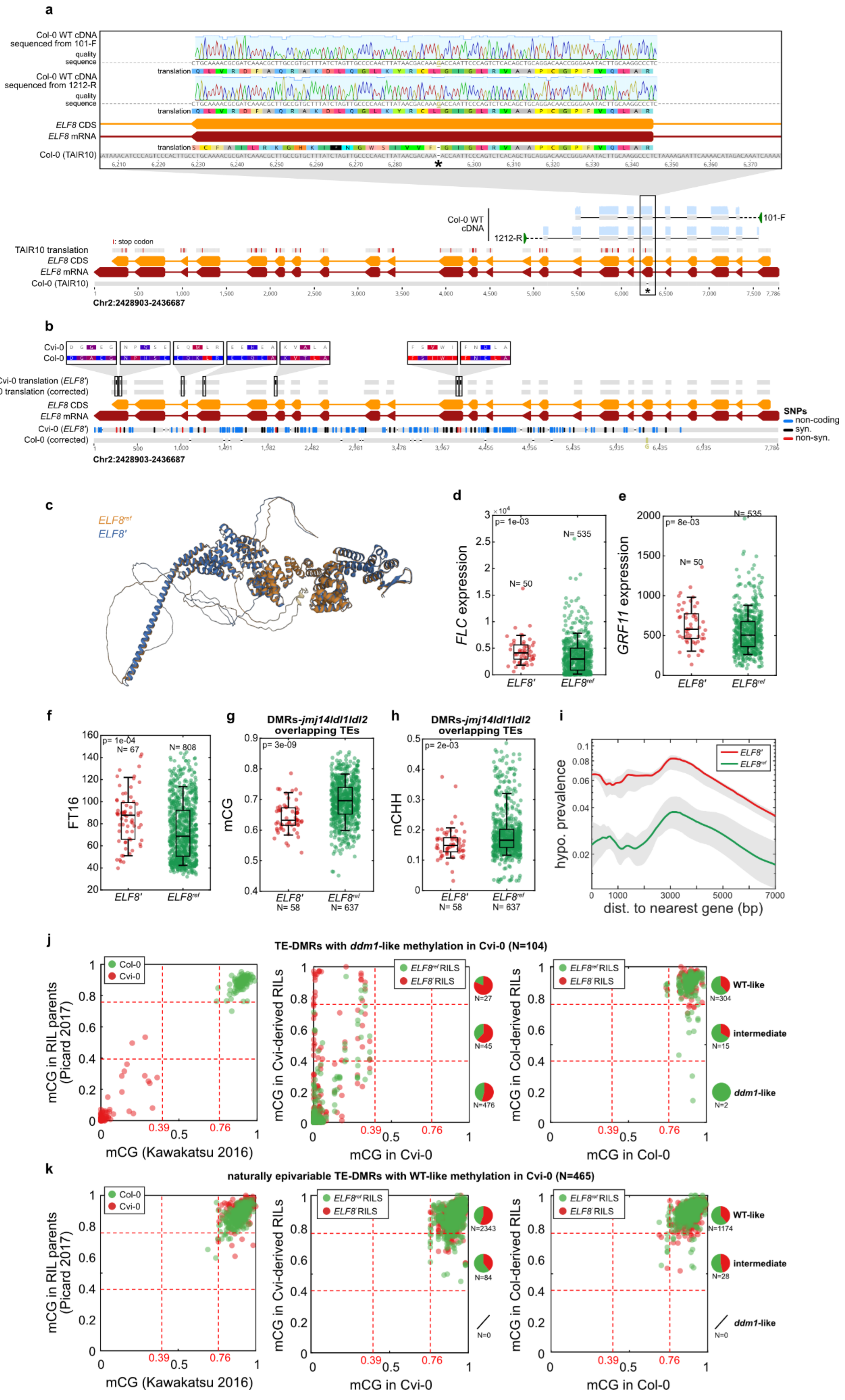

a
Col-0 WT cDNA sequenced from 101-F
quality
sequence
translation
Col-0 WT cDNA sequenced from 1212-R
quality
sequence
translation
ELF8 CDS
ELF8 mRNA
translation
Col-0 (TAIR10)
Col-0 WT cDNA
101-F
1212-R
I: stop codon
TAIR10 translation
ELF8 CDS
ELF8 mRNA
Col-0 (TAIR10)
Chr2:2428903-2436687
b
Cvi-0
Col-0
Cvi-0 translation (ELF8')
Col-0 translation (corrected)
ELF8 CDS
ELF8 mRNA
Cvi-0 (ELF8')
Col-0 (corrected)
Chr2:2428903-2436687
SNPs
non-coding
syn.
non-syn.
c
ELF8ref
ELF8'
d
p= 1e-03
N= 50
N= 535
FLC expression
ELF8'
ELF8ref
e
p= 8e-03
N= 50
N= 535
GRF11 expression
ELF8'
ELF8ref
f
p= 1e-04
N= 67
N= 808
FT16
ELF8'
ELF8ref
g
DMRs-jmj14ldl1ldl2 overlapping TEs
p= 3e-09
mCG
ELF8' N= 58
ELF8ref N= 637
h
DMRs-jmj14ldl1ldl2 overlapping TEs
p= 2e-03
mCHH
ELF8' N= 58
ELF8ref N= 637
i
ELF8'
ELF8ref
hypo. prevalence
dist. to nearest gene (bp)
j
TE-DMRs with ddm1-like methylation in Cvi-0 (N=104)
Col-0
Cvi-0
mCG in RIL parents (Picard 2017)
mCG (Kawakatsu 2016)
ELF8ref RILS
ELF8' RILS
mCG in Cvi-derived RILs
mCG in Cvi-0
N=27
N=45
N=476
mCG in Col-derived RILs
mCG in Col-0
WT-like N=304
intermediate N=15
ddm1-like N=2
k
naturally epivariable TE-DMRs with WT-like methylation in Cvi-0 (N=465)
Col-0
Cvi-0
mCG in RIL parents (Picard 2017)
mCG (Kawakatsu 2016)
ELF8ref RILS
ELF8' RILS
mCG in Cvi-derived RILs
mCG in Cvi-0
N=2343
N=84
N=0
mCG in Col-derived RILs
mCG in Col-0
WT-like N=1174
intermediate N=28
ddm1-like N=0

**Figure S10.**
(a) Sequence alignment of Col-0 WT *ELF8* mRNA sequenced using primers 101-F and 1212-R (indicated in green triangles) in comparison with predicted translation from TAIR10 sequence (stop codons are indicated in red). Top inset offers a detailed view of alignment over exon 6 where frame-shifting single-nucleotide deletion is predicted in the TAIR10 sequence (indicated by a *). (b) Sequence alignment of *ELF8* alleles of Col-0 (corrected, see Supplementary Note 6) and Cvi-0 (*ELF8'*) and comparison of predicted protein sequences. The 7 amino-acid substitutions between the two alleles are detailed with amino-acids color-coded by hydrophobicity (red: hydrophobic, blue: hydrophilic). (c) Alignment of protein structures of *ELF8* alleles of Col-0 (orange) and Cvi-0 (blue) predicted using AlphaFold (d,e) Comparison of *FLC* and *GRF11* expression levels between strains carrying the derived or the reference *ELF8* allele. (f) Comparison of flowering time at 16°C (FT16) between strains carrying the derived or the reference *ELF8* allele. (g,h) mCG and mCHH levels over DMRs identified in the *jmj14 ldl1 ldl2* triple mutant and that overlap TEs within strains carrying the *ELF8'* or $ELF8^{ref}$ allele. (i) Prevalence of natural epivariation in relation to the distance to the nearest gene among 50 randomly sampled strains carrying $ELF8^{ref}$ or *ELF8'* (10 replicates, max and min lowess values in shades of gray). (j) mCG levels in Col-0 *ddm1.2*, Col-0, and Cvi-0, as well as in 10 Cvi-0 x Col-0 RILs (102) depending on their genotype at the *ELF8* locus, over natural Cvi-0 epivariant TE-DMRs. Distributions of each genotype ($ELF8^{ref}$ in green, *ELF8'* in red) within the three methylation states (WT-like, intermediate, and *ddm1*-like) are represented as pie charts. (k) mCG levels in Col-0 *ddm1.2*, Col-0, and Cvi-0, as well as in 10 Cvi-0 x Col-0 RILs (102) depending on their genotype at the *ELF8* locus, over TE-DMRs with WT-like methylation in the Col-0 replicates and that are within 2kb (left panel) or further (right panel) from the nearest gene. Distributions of each genotype ($ELF8^{ref}$ in green, *ELF8'* in red) within the three methylation states (WT-like, intermediate, and *ddm1*-like) are represented as pie charts. Note that the pericentromeric regions of chr5 are derived from Cvi-0 in the 10 RILs analyzed, which explain the higher number of TE-DMRs inherited from that parent than from Col-0.

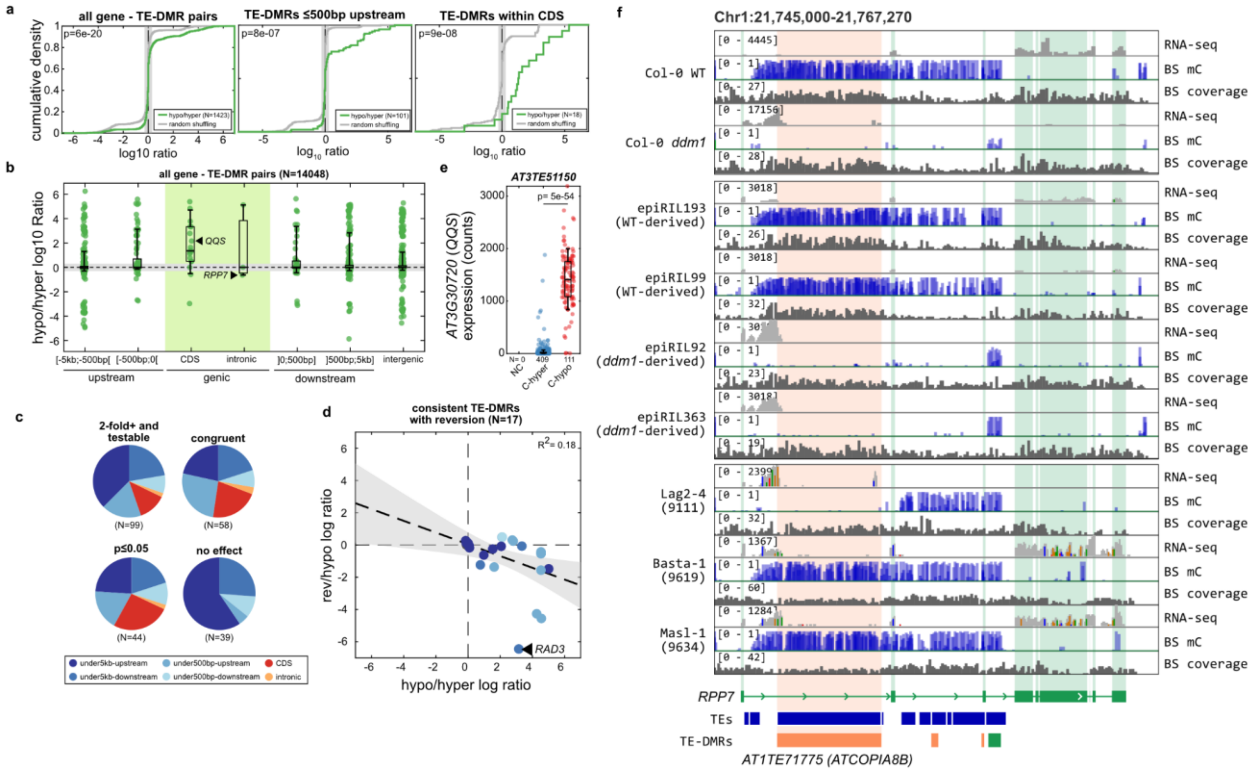


**Figure S11.**
(a) Distribution of expression differences of all genes bordering or in the immediate vicinity (≤500bp upstream or within the CDS) of naturally epivariable TE-DMRs between strains with *ddm1*-like vs WT-like methylation (hypo/hyper $\log_{10}$ ratio in green). The distribution of $\log_{10}$ ratios when the *ddm1*- or WT-like labels are randomly shuffled 10 times among all strains with RNA-seq data is represented in gray. The vertical shaded area corresponds to expression changes <2-fold in either direction. The p-value of the two-sample Kolmogorov-Smirnov test between the hypo/hyper and the random shuffled $\log_{10}$ ratio values is indicated. (b) log10 ratios by localization of each focal TE-DMR in relation to the nearby gene. The horizontal shaded area corresponds to expression changes <2-fold in either direction. (c) Position of TE-DMRs relative to the nearest gene for all 99 naturally epivariable TE-DMRs (left pie chart). The three other subsets shown (from left to right) contain naturally epivariable TE-DMRs with respectively congruent gene expression changes between these two settings, congruent gene expression changes that are also statistically significant in the epiRILs, no gene expression changes in the epiRILs. (d) Comparison of gene expression changes associated with reversion in the epiRILs (rev/hypo $\log_{10}$ ratio vs hypo/hyper $\log_{10}$ ratio) for each of the 17 TE-DMRs with relevant RNAseq data. Each dot is colored according to the location of the TE-DMR relative to the gene (same legend as in S11c). (e) Comparison of *QQS* expression levels between strains with or without epivariation at the fixed *QQS* TE-DMRs (NC=0). (f) BS-seq mC levels and coverage and RNA-seq read depth over *RPP7* TE-DMR at *ATCOPIA8B* (*AT1TE71775*) in the WT and *ddm1* parents, four epihaplotypically contrasted epiRILs and three natural strains, one of which (Lag2-4) exhibits epivariation over the TE-DMR.

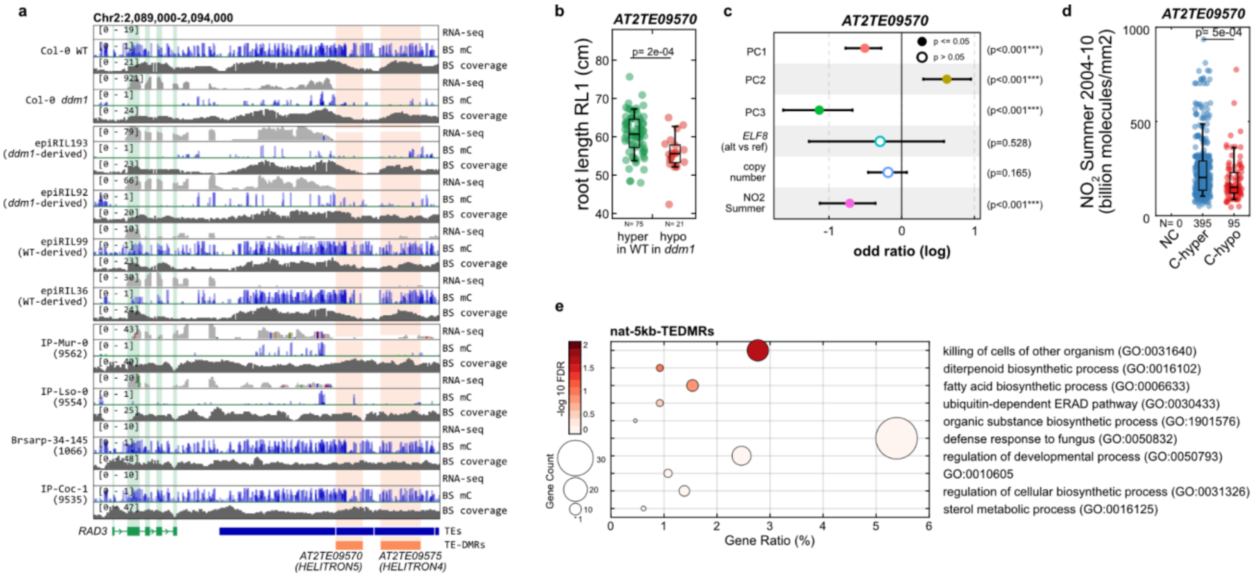

**Figure S12.**
(a) BS-seq mC levels and coverage and RNA-seq read depth over *RAD3* TE-DMRs (*AT2TE09570* and *AT2TE09575*) in the WT and *ddm1* parents, four epihaplotypically contrasted epiRILs and four strains, two of which exhibit epivariation over the two TE-DMRs. (b) Comparison of root length between epiRILs [(27)](#) in relation to the epiallelic state of the *RAD3* TE-DMR. (c) Marginal effects (log odd ratio) and p-values of population structure (three first PCs of kinship matrix), genotype at *ELF8*, copy-number of the TE family, and $NO_2$ atmospheric levels in summer (2004-2010) in GLM of natural epivariation occurrence at *RAD3* TE-DMR. (d) Comparison of $NO_2$ atmospheric levels in summer (2004-2010) at the collection site of strains with or without epivariation at the fixed *RAD3* TE-DMR. (d) Gene-Ontology enrichments (DAVID) of adjacent genes within 5kb of naturally epivariable TE-DMRs.

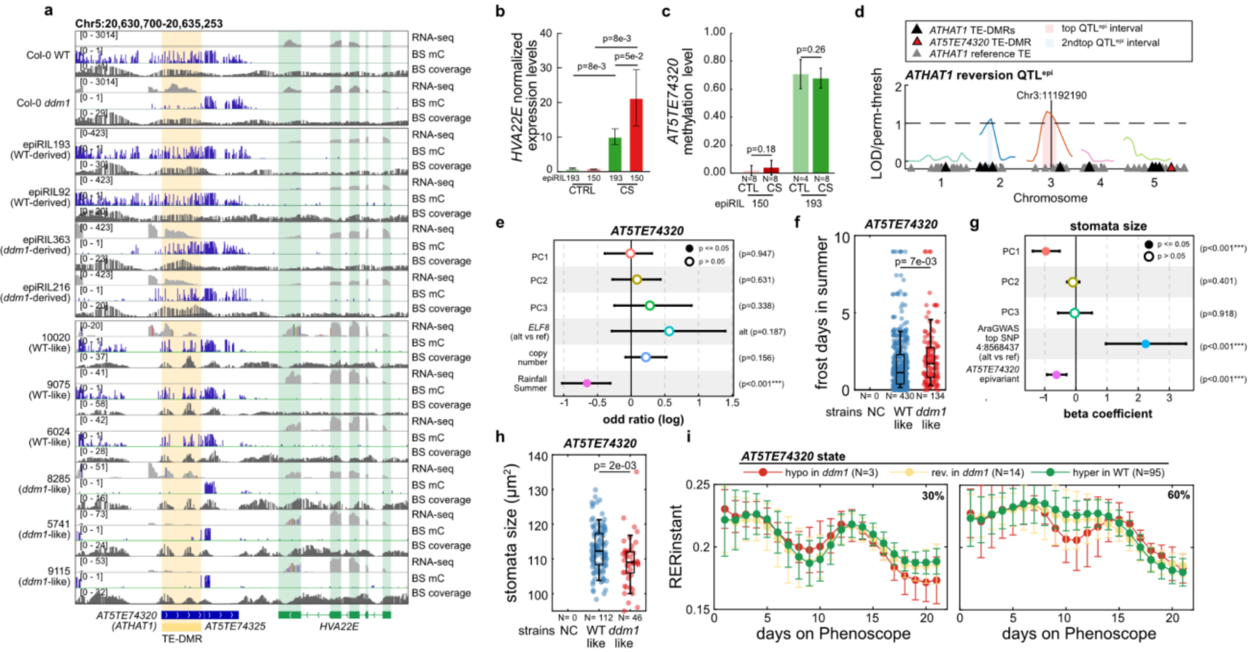


**Figure S13.**
(a) BS-seq mC levels and coverage and RNA-seq read depth over *HVA22E* TE-DMR at *AT5TE74320* in the WT and *ddm1* parents, four epihaplotypically contrasted epiRILs and four strains, two of which exhibit epivariation over the two TE-DMRs. (b) Expression level of *HVA22E* measured in three pools of 8 F10 seedlings grown in control conditions or exposed to cold-stress of epiRIL150, where the *HVA22E* TE-DMR is derived from *ddm1* and still hypomethylated inherited and of epiRIL193, where it is inherited from the WT parental line. (c) DNA methylation levels at *AT5TE74320* TE-DMR measured in 4-8 F10 individuals of the epiRIL150 and epiRIL193 grown under control conditions or exposed to cold-stress for 24h. (d) Interval mapping (LOD normalized by permutation threshold) of the reversion of all *ATHAT1* non-systematically reverting TE-DMRs in the epiRILs (cf Fig. 2,S5). The $QTL^{epi}$ intervals with significant LOD scores are highlighted. (e) Marginal effects (log odd ratio) and p-values of population structure (three first PCs of kinship matrix), genotype at *ELF8*, copy-number of the TE family, and rainfall in summer (2001-2010) in GLM of natural epivariation occurrence at *HVA22E* TE-DMR. (f) Comparison of number of frost days in summer at the collection site of strains with or without epivariation at the fixed TE-DMR (NC=0) downstream of *HVA22E*. The presence of outliers and their numbers are indicated by a triangle of the corresponding color. (g) Coefficients (beta) and p-values in GLM of stomata size (AraPheno 750) of population structure (three first PCs of kinship matrix), genotype at top SNP identified for this phenotype in AraGWAS (4:8568437), and natural epivariation at *HVA22E* TE-DMR. (h) Comparison of stomata size of strains with or without epivariation at the fixed TE-DMR (NC=0) downstream of *HVA22E*. (i) Daily growth rate (RER) across epiRILs grown on the Phenoscope in mild-drought (30% soil water content, top) or in well-watered conditions (60% soil water content, bottom) depending on the epiallelic state of the *HVA22E* TE-DMR.

**Table S1**

List of 7,023 TE-DMRs

**Table S2**

List of primers used for MSRE-detection of reversion in siblings of F9 epiRILs

**Table S3**

List of TE-DMRs targeted for single-molecule DNA methylation analysis in epiRIL238

**Table S4**

List of TE-DMRs with rare *ddm1*-like hypomethylation within WT-derived intervals

**Table S5**

List of 1,068 naturally epivariable TE-DMRs

**Table S6**

Statistics of 20 de novo assembled ONT genomes

**Table S7**

List of TE-DMRs hypomethylated in Cvi-0

**Table S8**

List of primers used for McrBC qPCR and methylation levels in Col WT and Cvi replicates

**Table S9**

34 TE-DMRs analyzed by McrBC-qPCR across 36 Cvi-0 x Col-0 RILs

**Table S10**

List of 99 TE-DMRs compared for nearby gene expression changes in the epiRILs and nature

**Table S11**

List of primers used for RT-PCR and RT-qPCR

**Dataset S1**

Details of DMRs by inheritance class

**Dataset S2**

List of 7,151 Mendelian TE-DMRs with at least 5DMSs covered in all 120 epiRILs

**Dataset S3**

Epihaplotypes at 1092 marker DMRs

**Dataset S4**

Methylation state of 7,023 TE-DMRs across 120 F9 epiRILs

**Dataset S5**

Methylation state of 7023 TE-DMRs across 720 natural strains

**Dataset S6**

Statistics of comparisons between BS-seq reads mapped on TAIR10 vs on 20 de novo assembled ONT genomes

**Dataset S7**
Growth rate measurements of epiRILs on Phenoscope in mild drought (0.3 SWC) and well-watered (0.6 SWC) conditions. Outliers removed from analysis are noted with a 1.
